# Reconstitution of SPO11-dependent double-strand break formation

**DOI:** 10.1101/2024.11.20.624382

**Authors:** Zhi Zheng, Lyuqin Zheng, Meret Arter, Kaixian Liu, Shintaro Yamada, David Ontoso, Soonjoung Kim, Scott Keeney

## Abstract

Homologous meiotic recombination starts with DNA double-strand breaks (DSBs) generated by SPO11 protein^1^. SPO11 is critical for meiosis in most species but the DSBs it makes are also dangerous because of their mutagenic^2^ and gametocidal^3^ potential, so cells must foster SPO11’s beneficial functions while minimizing its risks^4^. SPO11 mechanism and regulation remain poorly understood. Here we report reconstitution of DNA cleavage in vitro with purified recombinant mouse SPO11 bound to its essential partner TOP6BL. Similar to their yeast orthologs^5,6^, SPO11– TOP6BL complexes are monomeric (1:1) in solution and bind tightly to DNA. Unlike in yeast, however, dimeric (2:2) assemblies of mouse SPO11–TOP6BL cleave DNA to form covalent 5′ attachments requiring SPO11 active site residues, divalent metal ions, and SPO11 dimerization. Surprisingly, SPO11 can also manifest topoisomerase activity by relaxing supercoils and resealing DNA that it has nicked. Structure modeling with AlphaFold3^7^ illuminates the protein-DNA interface and suggests that DNA is bent prior to cleavage. Deep sequencing of in vitro cleavage products reveals a rotationally symmetric base composition bias that partially explains DSB site preferences in vivo. Cleavage is inefficient on complex DNA substrates, partly because SPO11 is readily trapped in DSB-incompetent (presumably monomeric) binding states that exchange slowly. However, cleavage is improved by using substrates that favor DSB-competent dimer assembly, or by fusing SPO11 to an artificial dimerization module. Our results inform a model in which intrinsically feeble dimerization restrains SPO11 activity in vivo, making it exquisitely dependent on accessory proteins that focus and control DSB formation so that it happens only at the right time and the right places.

## Introduction

SPO11-generated DNA double-strand breaks (DSBs) are a nearly universal feature of meiosis because they initiate the homologous recombination that supports chromosome pairing, chromosome segregation, and genome diversification, which are fundamental to gamete formation and sexual reproduction^1,8,9^. In mice and humans, absence of SPO11 or its partner TOP6BL (Topoisomerase VIB-like) results in infertility because of meiotic failure^10-13^.

SPO11 is evolutionarily related to the DNA cleaving subunit of archaeal Topoisomerase VI^14,15^. The Topo VI holoenzyme is a tetramer of two A subunits (Top6A, DNA cleaving) and two B subunits (Top6B, GH-KL-type ATPase)^16,17^ **(Fig. 1a)**. Concerted attack by the Top6A subunits on opposite strands of a duplex using active-site tyrosines breaks the DNA and produces covalent 5′-phosphotyrosyl links^18^. A winged-helix (WH) domain in Top6A bears this tyrosine (equivalent to Y138 in mouse SPO11), and a separate Rossmann fold known as a Toprim domain coordinates metal ions also needed for catalysis^16,17,19^. DNA strand breakage requires that the tyrosine of one Top6A protomer interact with the Mg^2+^-binding pocket of the second Top6A, forming a hybrid active site and necessitating Top6A dimerization for catalysis **(Fig. 1a)**.

**Fig. 1.**
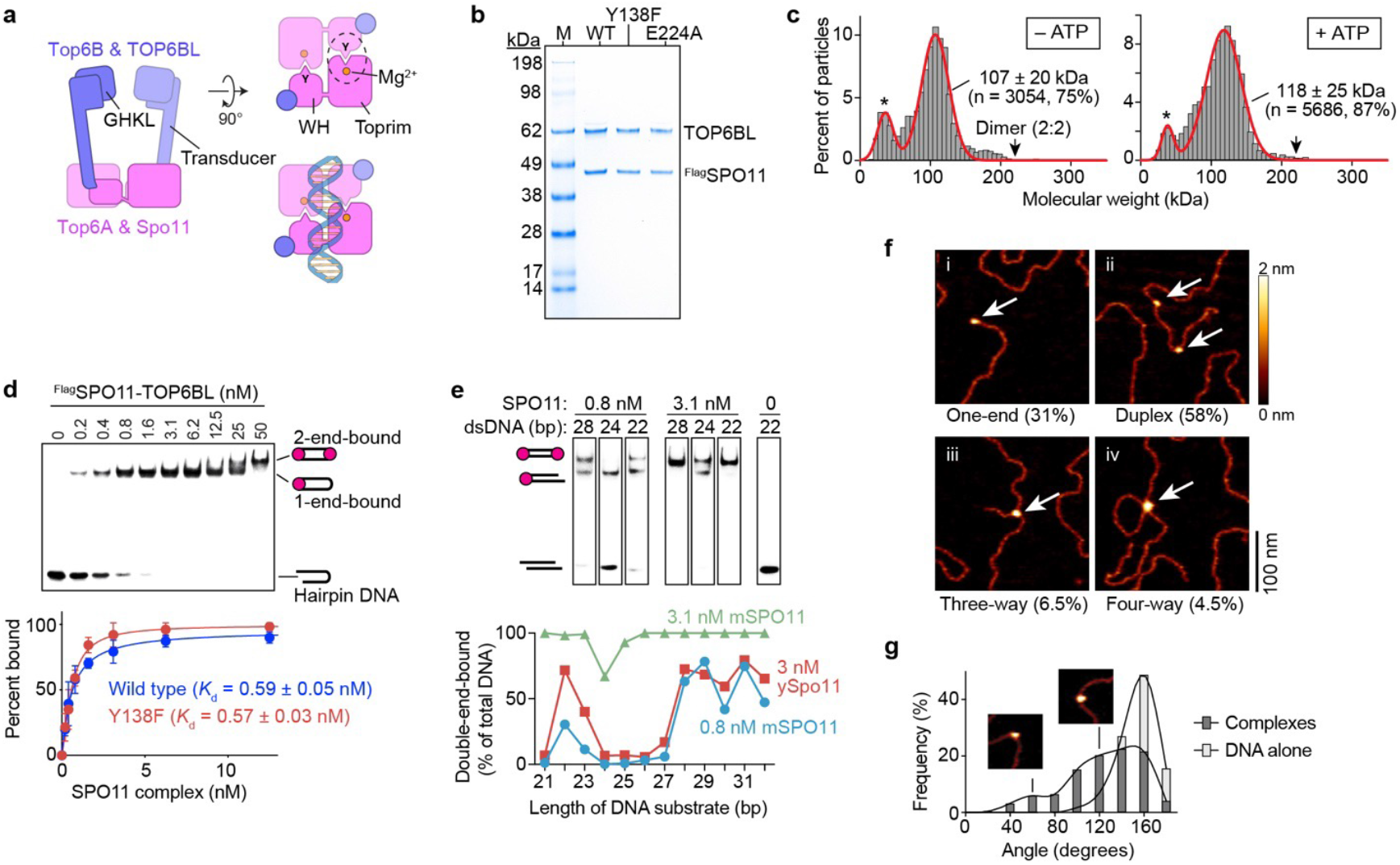
Purification and DNA-binding activity of SPO11–TOP6BL complexes. **a**, Domain organization of dimeric SPO11–TOP6BL complexes and Topo VI holoenzyme. Left, side view. Right, top views (with and without DNA) looking down into the DNA-binding channel. Catalytic tyrosine (Y), metal binding pocket, and hybrid active site (dashed circle) are shown. Reproduced from ref. ^5^ under a CC-BY 4.0 license. **b**, Coomassie-stained SDS-PAGE of purified ^Flag^SPO11–TOP6BL preparations (0.5 µg each). WT, wild type. **c**, Monomeric SPO11–TOP6BL complexes. Mass photometry profiles are shown without or with 5 mM ATP. Protein concentration was 28 nM. Particle counts (gray bars), gaussian density fits (red lines), fitted mean ± s.d., and percentages of total particles are shown. Asterisks, background material also present in blanks. **d**, EMSA of binding to DNA ends. SPO11 complexes were titrated with a 5′-labeled 25-bp hairpin substrate with a two-nucleotide 5′ overhang end. Quantification (mean ± s.d. of n = 3 experiments; apparent *K*_d_ given as mean ± s.e.) is shown below for wild type and Y138F (gel image in **Extended Data Fig. 1c**). **e**, DNA length dependence for double-end binding. Selected lanes from gel shift assays show binding of SPO11 complexes to DNAs of the indicated lengths with two-nucleotide 5′ overhangs on both ends (full gels in **Extended Data Fig. 1d**). Quantification of double-end binding is shown below, with yeast Spo11 data^32^ for comparison. **f**, AFM analysis of binding to linearized plasmid DNA. Examples are shown of binding to ends (one-end), internally on duplex DNA (duplex), junctions of three DNA arms (three-way), and junctions of four DNA arms (four-way). Percentages are from n = 200 particles scored. **g**, Histogram of bending angles (n = 200 particles). Examples are from subpopulations with modal values of ∼60° and ∼120°, similar to yeast^6^. Angles at randomly chosen positions along the DNA are shown as a control (n = 138 positions).

SPO11 covalently attaches itself to DNA during DSB formation in vivo^15,20^, indicating that it too cuts DNA by a topoisomerase-like transesterase reaction rather than a hydrolytic nuclease activity. However, despite the nearly three decades since SPO11 was recognized as the DNA-cleaving initiator of meiotic recombination^14,15^, key questions remain about its molecular mechanism because its activity has thus far been recalcitrant to reconstitution in vitro.

Top6B homologs were discovered more recently in plants, mice, yeasts and flies^11,21^. The homolog in *Saccharomyces cerevisiae*, Rec102, was long known to be important for Spo11 activity^22,23^, but its relationship with Top6B had gone unnoticed. The yeast and fly Top6B homologs lack the GHKL domain that mediates ATP-dependent dimerization in Topo VI, whereas the mouse and plant counterparts contain a degenerate version of this domain^11,21,24^.

Previous attempts to purify recombinant proteins often resulted in poor solubility after expression in *Escherichia coli*^25-28^. Purified plant SPO11 orthologs have moderate affinity for binding to duplex DNA (apparent *K*_d_ of 0.3 to 0.5 µM), but the only reported instance of DNA cleavage activity in vitro — for one of the five SPO11 homologs present in rice — did not involve covalent protein attachment to the DNA and was not shown to require the putative catalytic residues^26^. Mouse SPO11–TOP-6BL complexes were purified after expression in *E. coli*, but it was not clear if the material retained biologically relevant activity such as DNA binding or DNA cleavage^11^.

We recently reported the purification and characterization of recombinant *S. cerevisiae* Spo11 “core complexes” (Spo11 plus Top6B homolog Rec102 and the phylogenetically restricted interaction partners Ski8 and Rec104) after expression in insect cells^5,6^. Surprisingly, the yeast core complexes are “monomeric” in solution (1:1:1:1 stoichiometry) rather than having the expected 2:2:2:2 stoichiometry that is presumptively required for DSB formation and that is predicted from the stable dimerization of Top6A alone or within the Topo VI holoenzyme^16,17,19^. Monomeric core complexes bind with high affinity (sub-nanomolar *K*_d_) to DNA ends that mimic the cleavage product in having two-nucleotide (nt) 5′ overhangs, and core complexes (of unknown stoichiometry) also bind with lower but still high affinity internally to duplex DNA (*K*_*d*_ ∼ 5 nM)^6^. Cryo-EM structures revealed the subunit architecture and both protein-protein and protein-DNA interactions within DNA end-bound monomeric complexes that are thought to mimic the post-DSB state^5^. To date, however, no DNA cleavage activity has been reported.

Although many aspects of DSB formation are evolutionarily conserved, there are also key differences^29^. For example, there is no evidence for a role for Ski8 homologs outside of fungi^30^ and no clear orthologs of Rec104 outside of the family Saccharomycetaceae^5^. Moreover, the amino acid sequence, tertiary structure, and domain composition of the Top6B homologs in yeast and mouse differ strikingly^5,24^. These differences make it important to study SPO11 and its partners in different taxa, particularly in mammals. To this end, we report here the purification and characterization of recombinant mouse SPO11–TOP6BL complexes, reconstitution of DNA cleavage in vitro via a bona fide transesterase mechanism requiring SPO11 dimer formation, and structural models of dimeric complexes bound to DNA. Our findings suggest that a weak dimer interface is a hallmark of SPO11 that restrains its potentially dangerous ability to damage the genome and renders SPO11 dependent on other factors to control when and where it acts.

## Results

### Purification of 1:1 SPO11–TOP6BL complexes

We obtained complexes of Flag-tagged mouse SPO11 with TOP-6BL by affinity purification and size exclusion chromatography (SEC) after expression in cultured human cells (**Fig. 1b and Extended Data**

**Fig. 1a,b**). The proteins co-eluted in SEC at a volume consistent with a 1:1 complex (calculated mass of 110.3 kDa) (**Extended Data Fig. 1b**). This “monomeric” configuration of the complex was confirmed by mass photometry, in which a large majority of the particles were close to the predicted size for a 1:1 complex and few if any were of the size expected for a 2:2 stoichiometry (**Fig. 1c, left**). (Throughout, we will use “monomeric” and “dimeric” to refer to the number of SPO11 protomers in a complex.) The complexes remained monomeric in the presence of ATP (**Fig. 1c, right**), similar to purified TOP6BL alone^31^. SPO11–TOP6BL complexes were suggested to form 2:2 complexes when expressed in *E. coli*, on the basis of low-resolution SEC and glycerol gradient sedimentation analyses^*11*^, but misfolding or nonspecific aggregation could not be excluded. Thus, at least under the conditions tested here, mouse SPO11 complexes are monomeric in solution, similar to yeast Spo11 core complexes^5,6^ and unlike Top6A alone or in complex with Top6B^16,17,19^.

### High-affinity binding to DNA ends

We found that SPO11 complexes display tight, noncovalent DNA-end binding similar to yeast Spo11 core complexes. In electrophoretic mobility shift assays (EMSAs) with 25-bp hairpin substrates with a 2-nt 5′ overhang, protein concentrations as low as 0.2 nM yielded a single discrete shifted species (**Fig. 1d**). This high affinity DNA binding (apparent *K*_d_ ∼ 0.6 nM) was not diminished by mutating the catalytic tyrosine to phenylalanine (**Fig. 1d and Extended Data Fig. 1c**). A second shifted species appeared at higher concentrations (≥25 nM), likely reflecting binding of a second protein complex to the hairpin end.

Similar to yeast core complexes^32^, two mouse SPO11 complexes can bind separately and with high affinity to each end of a DNA duplex if the two 5′ overhangs are separated by at least 28 bp (**Fig. 1e and Extended Data Fig. 1d**). At low protein concentration (0.8 nM), double-end binding was strongly disfavored for DNA duplexes of 24–27 bp, but could be partially restored on even shorter DNA (22–23 bp) (**Fig. 1e**). The cryo-EM structure of yeast core complexes indicates that this behavior reflects steric clashes that are relieved by rotating the two end-bound complexes relative to one another^5^. These steric constraints help to explain the minimum spacing between adjacent DSBs in vivo when multiple Spo11 complexes cut the same DNA molecule in yeast or mouse^5,32^. Different from the yeast proteins, however, increasing the mouse protein concentration to only a few fold over the apparent *K*_d_ largely or completely restored double-end binding to all of the short DNA substrates tested (**Fig. 1e and Extended Data Fig. 1d**). The mouse protein thus appears to be more permissive of binding to DNA in close quarters.

Atomic force microscopy (AFM) analysis of binding to linearized plasmid substrates showed that most of the protein particles were bound internally to DNA duplexes and were frequently associated with DNA bends (**Fig. 1f,g**). Additionally, nearly a third of particles were bound to DNA termini (**Fig. 1f.i**). This finding confirms that SPO11 complexes prefer end binding because potential internal binding sites are in molar excess over termini. A minor fraction of particles was associated with places where duplex segments crossed one another or at locations with three DNA arms emanating out, likely representing simultaneous binding to an end and internally (**Fig. 1f.iii and 1.f.iv**). These binding patterns are congruent with those of the yeast core complex^6^.

### Reconstituting DSB formation

To test for DNA cleavage, we incubated SPO11 complexes with supercoiled plasmid DNA in the presence of Mn^2+^. We expected no activity for the mouse proteins — like their yeast orthologs — because we did not include their essential cofactors MEI4, REC114, IHO1, and MEI1 (reviewed in ^1^). Remarkably, however, wild-type complexes gave robust DNA cleavage that was not seen with the catalytic site mutant SPO11-Y138F (**Fig. 2a**). Both nicks and DSBs were generated, with nicks initially predominating: ∼60% of broken plasmids at 1 min had sustained only a nick.

**Fig. 2.**
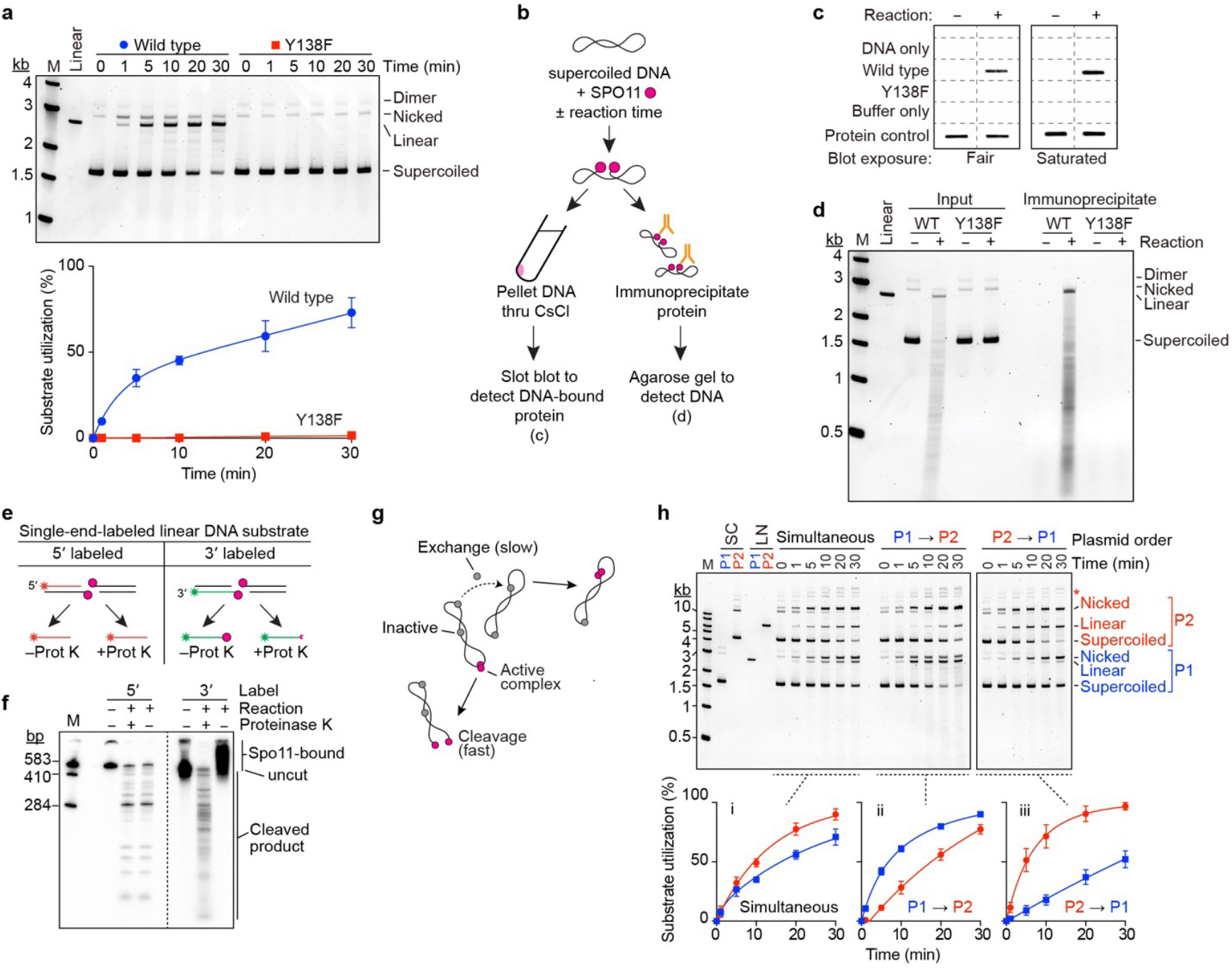
Reconstitution of SPO11-dependent DSB formation in vitro. **a**, DNA cleavage assays with SPO11–TOP6BL complexes containing wild-type or Y138F SPO11. Reactions contained 4 ng/µl pUC19 DNA, 100 nM SPO11 complexes, and 5 mM MnCl_2_. Deproteinized samples were separated on agarose gels stained with SYBR Gold. A representative gel is shown above, quantification (mean ± s.d. of n = 3 experiments) is below. **b-d**, Covalent attachment of SPO11 to cleaved DNA. Panel **b** shows a schematic overview of experiments in **c** and **d**. Wild-type or Y138F SPO11 complexes (325 nM) were mixed on ice with 4 ng/µl DNA and 5 mM MnCl_2_ in 60 µl and either immediately quenched with 0.5 % SDS (– reaction) or incubated at 37 °C for 11 min before quenching (+ reaction). In **c**, mixtures were centrifuged through CsCl cushions and the precipitated material was immuno-slot-blotted with anti-Flag antibodies (two exposure levels shown). No-protein negative controls (DNA only and buffer only) and positive controls for protein detection (10 ng) were included. In **d**, mixtures were immunoprecipitated with anti-Flag antibodies, then samples digested with proteinase K were separated by agarose gel electrophoresis. **e**, Differential prediction of 5′ covalent protein association with radiolabeled strands (asterisks) for 5′ vs. 3′ end-labeled substrates. **f**, Covalent attachment of SPO11 to 5′ ends. A 583 bp restriction fragment from pUC19 was either 5′ or 3′ radiolabeled on one end, then incubated with SPO11 complexes and separated by denaturing PAGE with or without prior digestion with proteinase K. **g**, Model to explain biphasic reaction kinetics. See text for details. **h**, Substrate order of addition determines reaction rate. SPO11 complexes (100 nM) were incubated with 4 ng/µl each of pUC19 (P1, 2.7 kb) and pCD-NA3.1-based plasmid mp134 (P2, 6.8 kb). Protein was mixed with both plasmids on ice before initiating reactions by transfer to 37 °C (simultaneous), or protein was incubated with one plasmid on ice and the second plasmid was added immediately before transfer to 37 °C (P1®P2 and P2®P1). Aliquots at the indicated times were quenched with SDS and deproteinated before agarose gel electrophoresis. Each plasmid (either supercoiled (SC) or linearized (LN)) was also run separately as a size marker. Representative gels are shown above, quantification (mean ± s.d. of n = 3 experiments) is below. Asterisk, slower migrating species that are likely to be multimers and/or catenated copies of P2.

Transesterase activity should leave SPO11 covalently attached to the DNA. To test this, we performed reciprocal experiments to look for stable SPO11-DNA association after cleavage (**Fig. 2b**). First, we purified the cleaved DNA by ultracentrifugation through a CsCl cushion that allowed DNA (plus covalently attached protein) to pellet, while free protein floated on top of the cushion. The pelleted DNA was dissolved, then ^Flag^SPO11 was detected by immunoblotting on a slot blot probed with anti-Flag antibody. Wild-type SPO11 copurified with the DNA as expected, but no copurifying protein was detected in negative controls that used SPO11-Y138F or that allowed SPO11 to bind but not react with the DNA (**Fig. 2c**).

In the second experiment, we immunoprecipitated SPO11, then assayed on agarose gels for coprecipitated DNA. Again as expected, broken DNA molecules were recovered in the anti-Flag immunoprecipitate from reactions with wild-type SPO11 protein, but not from reactions with SPO11-Y138F or from an unreacted control with wild-type protein (**Fig. 2d**).

Bona fide SPO11 activity should further leave the protein attached specifically to 5′ strand termini^33-35^. To test this, we digested linear DNA that was either 5′-or 3′-radiolabeled on one end. Radioactive cleavage products of the 5′-labeled substrate should not be protein-bound, and so should run identically on denaturing PAGE with or without protease digestion (**Fig. 2e, left**). Conversely, cleavage of the 3′-labeled substrate should leave SPO11 covalently attached to the radioactive strand, necessitating protease digestion for the labeled products to run into the gel (**Fig. 2e, right**). These predictions were met (**Fig. 2f**), definitively establishing that we have recapitulated the transesterase activity expected for SPO11.

DNA cleavage required divalent metal ions, with Mn^2+^ supporting substantially greater activity than Mg^2+^, which yielded more nicks relative to DSBs (**Extended Data Fig. 2a**). Ca^2+^ supported weak nicking but few if any DSBs (**Extended Data Fig. 2a**), in contrast with Topo VI from the hyperthermophilic archaeon *Saccharolobus shibatae*, for which Ca^2+^ supports robust DSB formation with little nicking^18,36^. DNA cleavage was greatest at physiological or higher temperature (**Extended Data Fig. 2b**) and showed a broad pH optimum from 7.5 to 8.5 (**Extended Data Fig. 2c**). Positively supercoiled and relaxed covalently closed plasmids were cut effectively, but activity was higher on negatively supercoiled DNA (**Extended Data Fig. 2d**,**e**). Linear DNA was also cut effectively (**Fig. 2f and Extended Data Fig. 2f**). Based on this optimization, we used standardized reaction conditions of 1 mM MnCl_2_, pH 7.5, and 37 °C for additional experiments unless indicated otherwise.

### Rapid DNA binding and slow exchange limit assembly of DSB-competent complexes

Reaction time courses typically displayed a fast initial rate of cleavage with little or no lag, followed by a longer phase of slower cleavage (e.g., **Fig. 2a and Extended Data Fig. 2d,e**). The slower phase is not an artifact from systematically underestimating total strand cleavage as substrates incur multiple nicks and DSBs, because it started at similar time points on different DNA substrates irrespective of the fraction of substrate cleaved (e.g., **Extended Data Fig. 2d,e**).

To account for these findings, we hypothesized that SPO11 complexes rapidly bind DNA when the reaction mixtures are assembled on ice, but only a subset of the protein-bound complexes are in a cleavage-competent state (possibly dimers) and are quickly consumed as they cut the DNA when shifted to 37 °C (**Fig. 2g**). We further posited that the sluggish second phase mostly reflects the exchange of non-productive SPO11-DNA complexes (possibly monomers) to form new cleavage-proficient complexes, which is slow relative to cleavage at least in part because of the high affinity (low off rate) of DNA binding and the excess of available monomer binding sites.

To test this hypothesis, we incubated SPO11 complexes with plasmid DNA on ice, then challenged the mixture with free DNA using a plasmid of a different size. In control experiments, both plasmids were cut with two-phase kinetics when mixed with the protein simultaneously (**Fig. 2h**, left). The larger plasmid was consumed more quickly as a percentage, as expected because it is a bigger target and thus fewer molecules were present in each reaction. In the staged reactions by contrast, the plasmid added first was cut substantially faster than either the plasmid added later or the same plasmid when in the simultaneous mixing regime (**Fig. 2h**, middle and right). Moreover, cleavage of the first plasmid again showed two-phase kinetics, whereas cleavage of the second plasmid followed a more uniform reaction time course that had an apparent rate comparable to the slow portion of the respective two-phase reaction. These findings support our interpretation (**Fig. 2g**).

### Computational model of a pre-DSB dimeric complex

To support the idea that SPO11 complexes can dimerize on DNA in a DSB-competent state, we used AlphaFold3^7^ to predict what such an assembly might look like. When queried with two copies of the SPO11– TOP6BL complex, a DNA duplex, and four Mg^2+^ ions, AlphaFold3 reproducibly predicted a dimeric protein assembly bound to bent DNA (**Fig. 3a–c, Extended Data Fig. 3a–d, and Supplemental Fig. 1, and Supplemental File 1**).

**Fig. 3.**
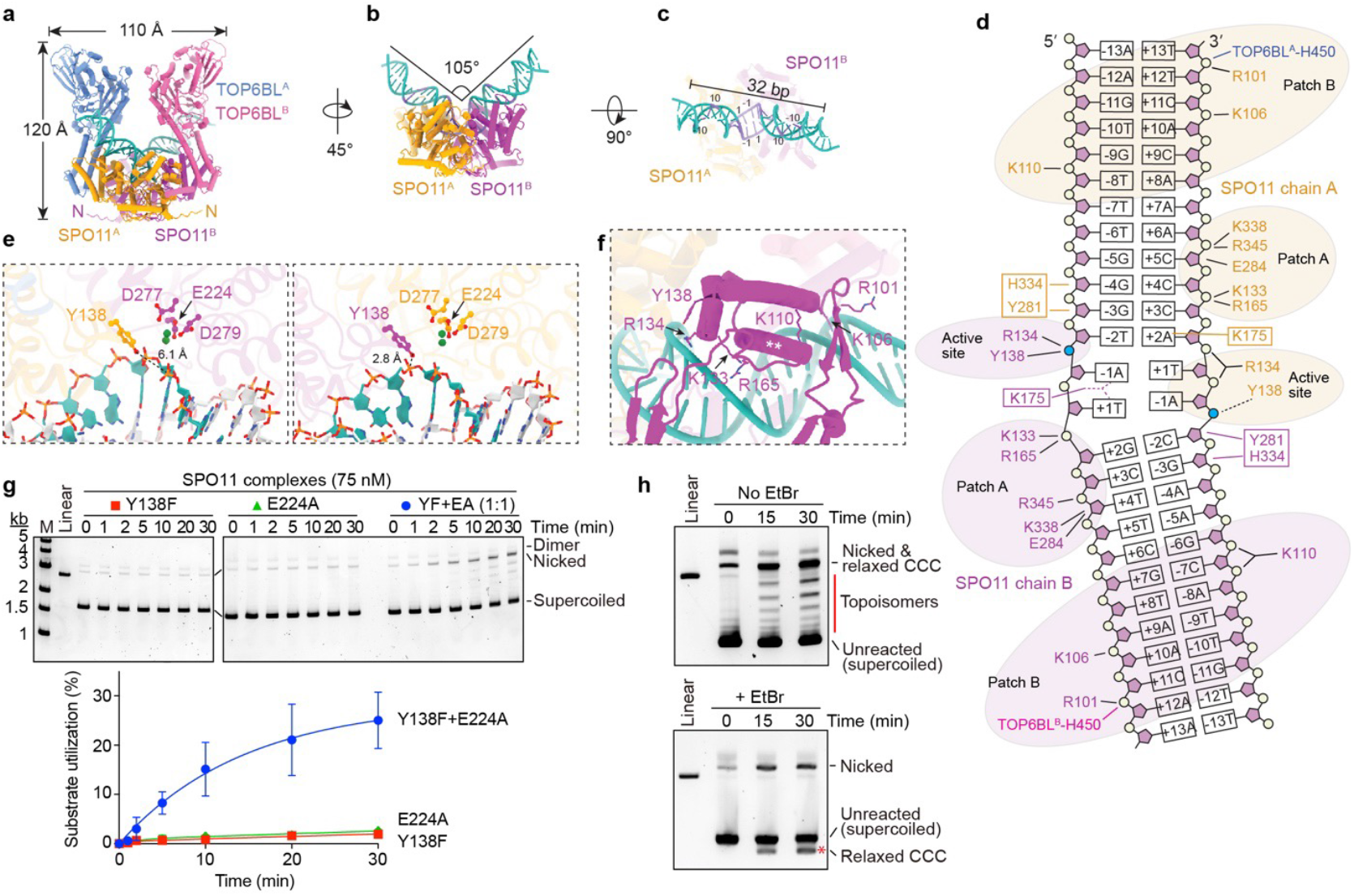
Assembly of DSB-competent dimeric complexes on DNA. **a–c**, AlphaFold3 model of a pre-DSB dimeric SPO11–TOP6BL complex on DNA. An overview of the entire structure (**a**), side view isolating SPO11 and the DNA (**b**), and a top view of the DNA (**c**) are shown. In **b** and **c**, DNA segments colored darker blue showed notable bias in base composition. In **c**, positions relative to dyad axis of the inferred cleavage site are numbered. A C-terminal α helix and long unstructured segment from TOP6BL^31^ are omitted for clarity (see **Extended Data Fig. 3a**). **d**, Inferred intermolecular contacts between the DNA (mostly sugar-phosphate backbone) and amino acid side chains of SPO11 and TOP6BL (both monomers of each) in an AlphaFold3 model. Groups of mostly positively charged residues (Patches A and B) and active site residues are shaded; boxed residues are predicted to occupy the minor groove. Bases are numbered by position relative to the presumed dyad axis of strand cleavage. **e**, Arrangement of catalytically important SPO11 residues in each of the two active sites (colored as in panel **a**). Distances of each Y138 residue to its cognate scissile phosphate are shown. **f**, Detail view of the positioning helix (**, residues 147–161) from a SPO11 WH domain, with adjacent DNA backbone contacts and the catalytic tyrosine indicated. **g**, Reconstitution of DNA nicking activity mixing catalytically defective SPO11 mutants. Representative agarose gels of cleavage reactions are above, quantification is below (mean ± s.d. of n = 3 experiments). **h**, Supercoil relaxation activity. Reactions containing mixtures of Y138F and E224A SPO11 complexes (as in **g**) were incubated for the indicated times then deproteinized and separated on agarose gels without (top) or with (bottom) ethidium bromide (EtBr). Red line highlights topoisomer ladder that disappears on EtBr-containing gels. Red asterisk highlights a new band on the EtBr gel that corresponds to plasmids that were relaxed covalently closed circles (CCC) after the reaction but that became positively supercoiled upon binding EtBr.

Supporting the model’s validity, it is congruent with empirical structures and functional data for homologous proteins^5,16,17,19,37,38^ (**Supplemental Discussion A**). For example, the secondary, tertiary, and quaternary protein architectures resemble crystal structures of Topo VI holoenzymes (**Extended Data Fig. 3e,f**). Moreover, the DNA is positioned along a channel in SPO11 cognate to the DNA binding surface defined experimentally for yeast Spo11 and proposed for Topo VI, and many of the protein-DNA contacts observed for yeast are recapitulated in the AlphaFold3 model (**Fig. 3a–d and Extended Data Fig. 3g–i**). Further, the model predicts the expected hybrid active sites with two Mg^2+^ ions positioned within each Toprim domain coordinated by side chains of E224, D277 and D279, and with the catalytic tyrosines nearby and close to opposite strands of the DNA (**Fig. 3e**), and the predicted SPO11 dimer interface includes similar structural elements as Top6A dimers (**Extended Data Fig. 4a**,**b**). Finally, DNA bending was observed in mouse (**Fig. 1g**) and yeast^6^ AFM experiments and in yeast cryo-EM structures^5^, and was predicted for Topo VI^38^. We conclude that the AlphaFold3 models are plausible representations of dimeric, DNA-bound, pre-DSB complexes.

**Fig. 4.**
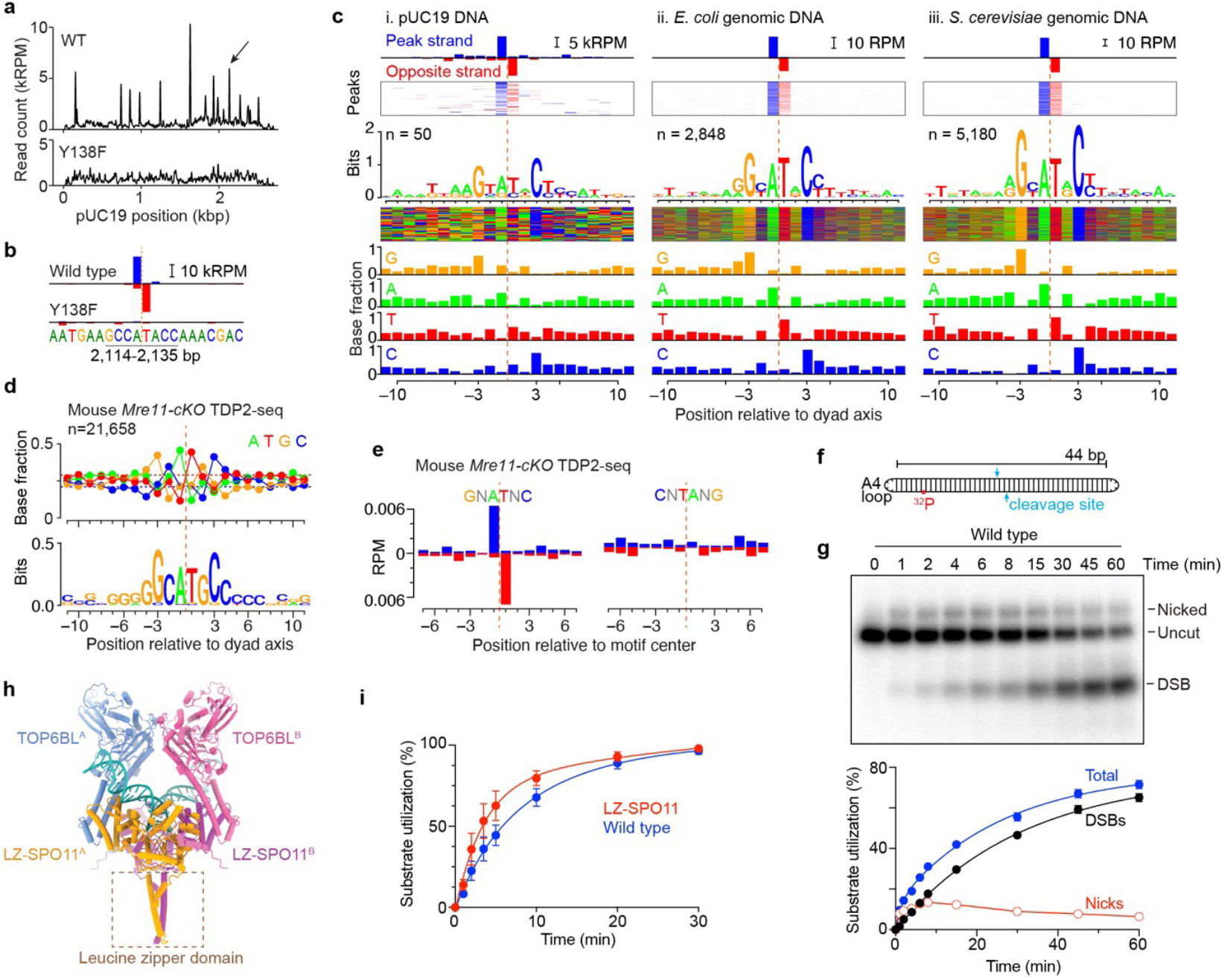
SPO11 sequence preferences and stimulation of cleavage by enhanced dimerization. **a**,**b**, TDP2-seq of in vitro cleavage reactions on pUC19 DNA with wild-type (WT) or Y138F mutant SPO11–TOP6BL complexes. Arrow in the pUC19 overview map (**a**) indicates the preferred cleavage position shown in detail view in **b**. Sequence read count is given in thousands of reads per million reads mapped (kRPM). Orange dashed line in **b** indicates dyad axis of cleavage. Averaged read count of two replicates is shown. The profile in **a** is smoothed with a 21-bp Hann filter. **c**, Base composition biases around preferred in vitro cleavage sites for reactions using pUC19 (**i**) or genomic DNA from *E. coli* (**ii**) or yeast (**iii**) as the substrate. The top two graphs show the strand-specific read counts at the indicated number of called peaks, either averaged (top) or as a heat map (second graph) where each horizontal line is one peak. The third graph shows the sequence logo. The fourth is a sequence color map where each horizontal line represents the sequence around one peak. The remaining bar graphs show the fractional composition of each base at each position relative to the dyad axis. **d**, Base composition bias for mouse SPO11 in vivo. Cleavage sites were identified within previously defined meiotic DSB hotspots^47^ in TDP2-seq maps from *Mre11* conditional knockout (cKO) mice, in which DSBs remain unresected^43^. Fractional base composition (above) and sequence logo (below) are shown. Note the different vertical scales for base composition and sequence logo compared to panel **c**, indicating weaker bias for the in vivo data. For **d** and **e**, representative data from one of two experiments are shown. **e**, Preferential DSB formation in vivo at positions matching the base composition that is preferred in vitro. Sites with sequence matching the in vitro bias (GNATNC, n = 11,423) or matching the non-preferred reverse sequence (CNTANG, n = 9,018) were identified within meiotic DSB hotspots. Average strand-specific TDP2-seq maps from *Mre11-cKO* mice are shown for each group of sites. **f**,**g**, Efficient cleavage of an oligonucleotide substrate containing a preferred cleavage site. Panel **f** shows a schematic of the substrate which was generated by self-annealing and ligation of a 5′ ^32^P-labeled ssDNA oligonucleotide containing the preferred cleavage site (blue arrows) from pUC19 shown in panel **b**. Panel g shows results from cleavage reactions with 10 nM wild-type SPO11 complexes and 0.5 nM oligonucleotide substrate. An autoradiograph of a representative gel is shown above, quantification is below (mean ± s.d. of n = 3 experiments). **h**, AlphaFold3 model of a dimeric SPO11–TOP6BL complex bound to DNA, with a leucine zipper (LZ) segment fused to each SPO11 N-terminus. The position and orientation of the LZ domain were variable between AlphaFold3 models. **i**, Stimulation of DNA cleavage by artificial SPO11 dimerization. Quantification is shown for cleavage reactions containing 4 ng/µl pUC19 DNA and 75 nM wild-type or LZ-tagged SPO11–TOP6BL complexes (mean ± s.d. of n = 3 experiments; representative gel image in **Extended Data Fig. 6g**).

Several new insights emerge. First, we asked whether AlphaFold3 could explain why ATP did not support dimerization of SPO11– TOP6BL complexes (**Fig. 1c**), unlike Topo VI, for which ATP stabilizes interactions between Top6B GHKL domains to form a second dimer interface^16,37^. When ATP was included in a query using Top6B proteins from *Saccharolobus shibatae*, AlphaFold3 predicted nucleotide-mediated GHKL domain dimerization that matched well with a crystal structure (**Extended Data Fig. 4c**)^37^, but not for TOP6BL (**Extended Data Fig. 4d**). The TOP6BL GHKL domain was predicted to have the expected secondary structure topology and overall fold (**Extended Data Fig. 4e**), but a structure-informed alignment indicated that TOP6BL lacks a G1 box (**Extended Data Fig. 4f**), a conserved glycine-containing element that directly contacts ATP^39^.

Second, DNA bending is predicted to be accompanied by substantial unwinding of the two base pairs that flank the predicted dyad axis (**Fig. 3c**). If correct, we speculate that this deformation may help to position the scissile phosphates in the active sites and/or impart strain on the DNA backbone to promote catalysis or disfavor religation after a DSB. The bending and unwinding may also explain nonrandom base composition at favored cleavage sites (addressed below).

Third, the relative arrangement of the WH and Toprim domains in each SPO11 monomer does not match precisely with the yeast Spo11 cryo-EM structure, which is thought to represent a post-DSB product-bound state^5^. However, the two structures can be aligned well by a 13° rigid-body movement of the WH domain (**Extended Data Fig. 4g**), which is plausible given that the protein segment connecting them is thought to be a flexible linker^6,19^. This finding provides support for the idea that DSB formation is accompanied by a SPO11 conformational change that may influence reversibility of the cleavage reaction^5^.

Fourth, although protein-DNA contacts are similar in both broad strokes and details to those for monomeric yeast core complexes (**Supplemental Discussion A**), there are also new features that were not previously apparent. For example, an α helix in the WH domain (residues 148–159) lies across the major groove at the bend (**Fig. 3f**). The side chains do not make obvious DNA contacts, but the helix’s position suggests that it braces two separate backbone-interaction patches on either side of it: K133/R134/R165 plus the catalytic Y138 on one side, and R101/K106/K110 on the other side. This helix might thus be important for positioning the two patches, thereby stabilizing the bend and helping to position Y138 near the scissile phosphate.

Another surprising feature of the protein-DNA contacts is that they are not rotationally symmetric. The same residues in each SPO11 monomer contact the DNA, but one monomer is shifted by ∼1 bp relative to the other (**Fig. 3d and Extended Data Fig. 3g,h**). This places one of the catalytic tyrosines further away from the scissile phosphate than the other (**Fig. 3e**), so this model is not consistent with both active sites being catalytically competent at the same time. This asymmetry should be viewed cautiously because it remains to be experimentally validated, but it may help to explain why nicks are a prominent early product in cleavage reactions.

### Each DNA strand is cut by a hybrid active site in a SPO11 dimer

To bolster the structure modeling, we sought direct experimental proof that strand cleavage is carried out by hybrid active sites in a dimer. To do this, we asked whether DNA cleavage activity could be restored by cross-complementation of SPO11 proteins that were inactive on their own because of catalytic site mutations in either the WH domain (Y138F) or metal-binding pocket of the Toprim domain (E224A). Because wild-type SPO11 can nick the DNA, we reasoned that mutant heterodimers would also generate nicks because they would have one complete active site and one doubly mutated one.

The E224A mutant retained normal DNA end binding (**Extended Data Fig. 4h**), unlike its yeast counterpart (Spo11-E233A)^6^, but was nonetheless incapable of cutting DNA (**Fig. 3g**). In contrast, a mix of Y138F and E224A mutant proteins yielded nicking activity without appreciable DSB formation (**Fig. 3g**). This result provides compelling evidence for formation of hybrid active sites.

### SPO11 relaxes and religates DNA

At later reaction time points with mixtures of Y138F and E224A, a ladder of bands appeared with intermediate electrophoretic mobility between supercoiled and nicked DNA (**Fig. 3h**). These bands disappeared when the agarose gels contained ethidium bromide, coincident with appearance of a new band running faster than the unreacted substrate (asterisk in **Fig. 3h, bottom**). This behavior is as expected for partially relaxed, covalently closed topoisomers that become positively supercoiled upon binding ethidium bromide, and rules out that these might be linear species from double cutting at defined sites in the plasmid.

We conclude that SPO11 complexes can exhibit topoisomerase activity to relax supercoiled DNA and that SPO11 can reseal a broken DNA strand(s) by reversing the covalent 5′-phosphotyrosyl linkage. Topoisomer ladders were also generated by wild-type protein: they were most prominent when nicking occurred without substantial double-strand cleavage (e.g., in the presence of Ca^2+^; **Extended Data Fig. 2a**), but could be observed whenever nicked circles remained present with both negatively and positively supercoiled DNA (**Extended Data Fig. 2d**). Thus, relaxation and resealing can occur on substrates that have been nicked, but whether religation can also occur after a DSB is not clear. A potential mechanism and implications of the supercoil relaxation activity are provided in **Supplemental Discussion B** and **Extended Data Fig. 4i**.

### Biased base composition at preferred cleavage sites

Preferred cleavage sites have been noted for yeast Spo11 in vivo^33,35,40,41^ and were apparent for the mouse protein in vitro (**Fig. 2f**). To delineate SPO11 sequence preferences, we mapped DSB positions by removing covalently bound protein with proteinase K and tyrosyl phosphodiesterase 2 (TDP2), blunting DNA ends with T4 DNA polymerase, ligating adaptors to the blunted ends, and deep sequencing the ligation junctions^42^ (**Extended Data Fig. 5a**).

With wild-type protein and pUC19 as the substrate, we recovered sequencing reads distributed across the plasmid but with prominent peaks that were not observed with SPO11-Y138F (**Fig. 4a**). Replicate maps were reproducible for wild type but not for the random background with SPO11-Y138F (**Extended Data Fig. 5b**). Cleavage products are expected to have a 2-nt 5′-overhang, the middle of which is a twofold rotational symmetry axis. Fill-in of the overhang by T4 DNA polymerase should yield top- and bottom-strand reads that overlap by 1 bp, giving an offset of +1 in a cross-correlation test between the strands (**Extended Data Fig. 5a**). Preferred cleavage sites matched this expectation for wild-type but not the SPO11-Y138F control (**Fig. 4b**,**c.i and Extended Data Fig. 5c**), demonstrating that the DSBs formed in vitro have the correct polarity.

Visual inspection of the ten strongest cleavage sites in pUC19 showed strong bias for a G at the –3 position relative to the dyad axis and C at the rotationally symmetric +3 position (**Extended Data Fig. 5d)**. To examine the sequence biases further, we averaged the base composition around 50 peaks in the sequencing maps. This analysis extended the strong signature of biased base composition beyond just the +3 and –3 positions, encompassing the 8 bp surrounding the dyad axis (**Fig. 4c.i)**. The sequence signature was rotationally symmetric, as expected for a pair of SPO11 complexes engaging a pair of half-sites on the DNA duplex. We emphasize that this is a base composition bias, not a strict sequence motif, and that the rotational symmetry does not imply that individual SPO11 cleavage sites are palindromic.

We also generated maps using *E. coli* or *S. cerevisiae* genomic DNA as the substrate. Prominent cleavage sites with the expected strand polarity were again observed with wild-type but not SPO11-Y138F protein (**Extended Data Fig. 5e**,**f**), with similar base composition biases (**Fig. 4c.ii and 4c.iii**). Similar results with both supercoiled plasmid and relaxed genomic DNA show that superhelical tension is not required to establish the bias. Importantly, however, the sequence signature correlated with the amount and complexity of the substrate: the signature was stronger with *E. coli* DNA than with pUC19 and stronger still with yeast DNA, allowing detection of sequence bias out to the –10/+10 positions.

We attribute the stronger sequence signatures to two factors. First, the scaled-up reactions with yeast or *E. coli* DNA had threefold higher protein concentration but ∼ninefold higher DNA concentration than the pUC19 reactions. The increased DNA:protein ratio is expected to attenuate dimer formation by favoring more dispersed binding of monomeric protein complexes. Second, every position on the DNA substrate competes with every other position for binding and cleavage by SPO11 complexes. The more complex the substrate, the lower the concentration of any given site and the higher the concentration of competing sites. We suggest that these factors together affect the stringency of site selection, making SPO11 more dependent on an optimal match to its preferred target site the more complex the substrate.

We further tested whether the same sequence signature applies in vivo. To do this, we performed TDP2-seq on *Mre11*-deficient mouse spermatocytes, which accumulate unresected DSBs^43^. Averaging over cleavage positions showed a weaker but similar base composition from –10 to +10 as in vitro, including enrichment for G/C at –3/+3 (**Fig. 4d**). Moreover, averaging over sites within known DSB hotspots that match a favored sequence (GNATNC) enriched for top- and bottom-strand reads at the correct positions, not seen with an unfavorable sequence (**Fig. 4e**). We conclude that the intrinsic substrate preferences of SPO11 in vitro also shape the DSB landscape in vivo, but to a weaker extent, presumably because additional factors also contribute (Discussion).

Not surprisingly, the biased sequence composition spans the predicted footprint of dimeric SPO11 complexes on the DNA (**Fig. 3c**,**d**). The strong bias for A and T at –1 and +1 corresponds to the position where the DNA is most distorted in the AlphaFold3 model, and SPO11 is predicted to contact bases near the dyad axis and to make multiple contacts to the DNA backbone spanning the biased central region. Notable interactions in light of the sequence signature include Y281 and H334 contacting one strand between –2 and –4 and Patch A contacting the opposite strand nearby, and Patch B contacting both strands near the –10 and +10 positions (**Fig. 3d** and **Extended Data Fig. 3g**).

What is driving the sequence bias is not currently clear. It is unlikely to reflect direct readout of the bases because nearly all of the protein contacts are with the DNA backbone. Instead, the preferred base composition may favor an intrinsic DNA shape that supports SPO11 activity. In this vein, the preferred base composition predicts systematic variation in minor groove width (**Extended Data Fig. 5g**). The local sequence composition may influence initial SPO11 binding, DNA bending, and/ or catalysis.

Interestingly, even though the mouse and yeast proteins have similarly high affinity for DNA ends, have similarly sized footprints on DNA, and share many of their DNA-binding residues in common, their sequence preferences differ markedly. Deduced from in vivo patterns^32,40,42,44^, yeast Spo11 strongly prefers A at –4 and –5 and C at –2 (the base immediately upstream of the scissile phosphate), but cares little about the –3 position or the dinucleotide centered on the dyad axis (**Extended Data Fig. 5h**). Because the mouse sequence bias is weak where the yeast bias is strong, and strong where yeast’s is weak, it appears that fine-scale site selection by SPO11 is evolutionarily plastic.

### Fostering dimerization improves cleavage activity

Armed with knowledge of SPO11 site preferences, we revisited the hypothesis that inefficient dimer formation is what limits overall cleavage activity. We reasoned that a DNA substrate optimized for SPO11 dimer assembly should improve cleavage efficiency. To test this, we made an oligonucleotide substrate containing a preferred cleavage site from pUC19 in the middle (**Fig. 4b,f**). Both DNA ends were A_4_ hairpins to reduce SPO11 end-binding affinity, and the duplex arm lengths (22 bp each) were chosen to encourage only a single dimeric complex to bind near the middle. This substrate was bound with high affinity (apparent *K*_d_ of 1.8 nM) and doubly-bound DNA complexes were efficiently formed (**Extended Data Fig. 6a)**.

As predicted, this substrate was cut well when most of the DNA molecules were bound by two SPO11 complexes (**Fig. 4g**). Both nicks and DSBs were formed (**Fig. 4g**), the reaction required Y138 of SPO11 (**Extended Data Fig. 6b**), and Mn^2+^ was strongly preferred over Mg^2+^ (compare **Fig. 4g** with **Extended Data Fig. 6c**). A mixture of Y138F and E224A mutant proteins generated only nicks, as expected (**Extended Data Fig. 6d**).

The plasmid reactions were inefficient on a per-protein basis. For example, even though most of the substrate was cut by 30 min with 100 nM protein in **Fig. 2a**, this represented less than ∼300 fmol of strand breaks and thus only ∼4% of the SPO11 molecules. With the oligonucleotide substrate and wild-type protein, 72% of the DNA was cut by 60 min (**Fig. 4g**). The fraction of DNA expected to be dimer-bound under the simplistic assumption of two independent monomer binding sites, each with a *K*_d_ of 1.8 nM, is the same (72%). This suggests that nearly every SPO11 dimer is capable of making a strand break on the oligonucleotide substrate. We infer from this high apparent efficiency that most or all of the protein preparation is active if it is able to assemble a dimer on a cleavage-susceptible sequence. We also note that there was little or no indication from the EMSAs that an appreciable population of higher order complexes contained a SPO11 dimer plus two DNA duplexes. Thus, our findings do not provide support for binding of a second duplex being required for cleavage^6,38,44^, although we do not exclude that such binding can occur and might affect cleavage in other contexts.

Finally, we tested whether cleavage efficiency could be increased by artificially tethering two SPO11–TOP6BL complexes together. We leveraged the predicted arrangement of the SPO11 N-termini on one face of the dimeric assembly away from the DNA-binding surface (**Fig. 3a**). AlphaFold3 was used to design a dimerization module consisting of a 33 amino acid leucine zipper (LZ)-forming segment from yeast GCN4^45^ followed by a flexible gly-ser linker fused to the SPO11 N-terminus (**Fig. 4h**). LZ-SPO11–TOP6BL complexes eluted earlier in SEC at a position close to theoretical for a dimeric complex (2:2 heterotetramer, 230.4 kDa) (**Extended Data Fig. 6e**), and 64% of SPO11–TOP6BL particles were consistent with dimeric complexes in mass photometry experiments at low protein concentration (15 nM) (**Extended Data Fig. 6f**).

As predicted, LZ-SPO11 complexes showed enhanced activity in the standard plasmid cleavage assay (**Fig. 4i and Extended Data Fig. 6g**). The initial cleavage rate was ∼1.7-fold faster than with wild-type SPO11 and the reaction consumed more of the substrate in the initial burst, as expected if a larger fraction of DNA-bound SPO11 started out in an active dimeric form.

## Discussion

We have found that mouse SPO11–TOP6BL complexes are monomeric in solution but can dimerize on DNA and then carry out strand cleavage. Many other proteins are essential for SPO11 activity in vivo in many species^1^, so it has been unclear whether SPO11 complexes by themselves would be sufficient. Our results agree well with studies by the Claeys Bouuaert and Tong laboratories.

We consider the weak dimerization of SPO11–TOP6BL complexes to be a defining distinction from the homologous Topo VI enzymes. In this regard, there are two important differences: a low intrinsic affinity for SPO11 self-association compared to Top6A, and loss of ATP-driven dimerization by the GHKL domain. As elaborated below, we suggest that weak dimerization is key to understanding SPO11 mechanism and regulation.

Previous studies in yeast showed that evolutionarily conserved Spo11 accessory factors Rec114, Mei4, and Mer2^IHO1^ assemble cooperatively to form large nucleoprotein condensates that in turn recruit Spo11 core complexes^46^. This is proposed to establish chromosome axis-associated clusters of co-oriented SPO11 proteins that can bind and cut DNA on chromatin loops^6,32,46^ (**Extended Data Fig. 7a**). Clustering of SPO11 proteins gives a high efficiency of DSB formation per cluster, despite a low efficiency of DSB formation per SPO11 core complex.

Our findings suggest the following key additions to this model. In vivo, where the amount and complexity of the DNA target is high relative to the amount of available SPO11 protein, we propose that weak dimer formation plus high affinity DNA binding together establish a kinetic trap that favors dispersed binding of monomers to chromatin and disfavors formation of catalytically competent dimers. SPO11 thus becomes highly dependent on its accessory factors that increase the local SPO11 concentration through clustering and also prepay part of the entropic cost of dimerization by co-orienting SPO11 complexes. The overall result is that SPO11 is nearly exclusively active within the context of higher order chromosomal structures that also provide a means to regulate the timing, number, location, and spacing of DSBs. This arrangement provides an elegant solution to the challenge of how to ensure that SPO11 carries out its essential function while minimizing the risks that poorly controlled DSBs would pose.

A corollary is that SPO11 clustering should diminish the dependency of DSB formation on an optimal DNA sequence. This may help to explain why sequence signatures of preferred cleavage sites in vivo are weaker than in vitro. Moreover, the DSB frequency at a given genomic location is determined by many factors operating over different size scales (e.g., targeting by the PRDM9 histone methyltransferase; nucleosome occupancy; higher order chromosome structure; replication timing, etc.)^40,47-51^. These factors shape DSB locations independent of the intrinsic affinity of SPO11 for the DNA at the cut site, so they also weaken the contribution of SPO11 itself.

It is interesting that the mouse protein is thus far more permissive for DSB formation in vitro than yeast Spo11. We speculate that this difference may trace in part to differences in the intrinsic favorability of monomeric vs. dimeric DNA-bound complexes. AlphaFold3 models suggest that SPO11 from mice may adopt similar conformations as monomers and dimers (**Extended Data Fig. 7b**,**c**), whereas yeast Spo11 monomers may tend to adopt a conformation that differs significantly from the dimer conformation (**Extended Data Fig. 7d**). We speculate that the protein from mouse (and perhaps a number of other species; **Extended Data fig. 7e**) may be intrinsically more dimer-compatible. If so, it suggests that SPO11 dimerization potential is itself an evolutionarily variable, and thus tunable, property.

## Methods

### Mouse experiments

Mouse experiments were performed in accordance with US Office of Laboratory Animal Welfare regulations and were approved by the Memorial Sloan Kettering Cancer Center Institutional Animal Care and Use Committee. Mice were maintained on rodent chow with continuous access to food and water and were euthanized by CO_2_ asphyxiation for tissue harvest. Male germline-specific *Mre11-cKO* mice were generated by crossing *Mre11-flox*^52^ and *Ngn3-Cre*^53^ alleles as described elsewhere^43^.

### Recombinant protein expression and purification

The full-length optimized coding DNA for mouse SPO11 (Uniprot: Q9WTK8-1) with a Flag tag followed by a TEV cleavage site at the amino terminus, and untagged TOP6BL (Uniprot: J3QMY9-1) were cloned into the pCDNA3.1(+) vector (Invitrogen). The SPO11 construct encodes for the b splicing isoform, which includes exon 2 that is skipped in the a isoform. SPO11b is both necessary and sufficient for most DSB formation in vivo, and the exon 2-encoded sequence is required for robust interaction with TOP6BL^11,20,54-56^. Point mutants (Y138F, E224A) were generated using QuikChange mutagenesis. The sequence encoding the GCN4 leucine zipper and GS linker was codon-optimized, synthesized as gBlocks, and then cloned into the same vector as mouse SPO11, positioned after the Flag tag and before the TEV cleavage site.

FreeStyle™ 293-F cells (Invitrogen) were cultured in FreeStyle™ 293 expression medium (Gibco) under 5% CO_2_ in a Multitron-Pro shaker (Infors, 120 rpm) at 37 °C. When the cell density reached 1.2 × 10^6^ cells per ml, the plasmids were co-transfected as follows. For a one-liter cell culture, 1 mg of plasmids were pre-incubated with 2 mg of 40-kDa linear polyethylenimine (PEI) (Polysciences) in 25 ml of Opti-MEM™ (Gibco) for 25–30 min. Transfection was initiated by adding the mixture to the cell culture. About 16 hr after transfection, sodium butyrate was added to a final concentration of 10 mM. Transfected cells were cultured for an additional 32 hr (for a total of 48 hr) before harvesting.

For each batch of protein purification, one liter of transfected cells was harvested by centrifugation at 3100 g and resuspended in lysis buffer containing 25 mM HEPES-NaOH, pH 7.5, and 500 mM NaCl. The suspension was supplemented with a protease inhibitor cocktail, including 1 mM phenylmethylsulfonyl fluoride (PMSF) and 1× Halt™ Protease Inhibitor Cocktail (Thermo Scientific). Cells were lysed by sonication and centrifuged at 46,000 g for 60 min. The supernatant was applied to anti-Flag M2 affinity resin (Sigma) and allowed to flow through by gravity at 4 °C. The resin was rinsed three times with lysis buffer. Proteins were eluted with lysis buffer containing 200 μg/ml Flag peptide (Sigma). The eluent was then concentrated using a 10-kDa cut-off Centricon (Millipore) and further purified by SEC (Superdex 200 Increase 10/300 GL, GE Healthcare). Peak fractions were pooled and concentrated to 40 μl at a concentration of approximately 1.5 mg/ml. Aliquots were frozen in liquid nitrogen and stored at –80 °C. Protein concentration was determined by absorbance at 280 nm. For experiments comparing wild type with LZ-SPO11, matched protein concentrations were further confirmed by SDS-PAGE and Coomassie staining.

### Oligonucleotide substrates for DNA binding and cleavage assays

Oligonucleotide sequences are provided in **Supplemental Table 1**. Hairpin substrates with a two-nucleotide 5′-overhang end for EMSAs were assembled by self-annealing oligonucleotides that were 5′-end labeled with [γ-^32^P]-ATP (Perkin Elmer) and T4 polynucleotide kinase (NEB), followed by purification by native PAGE. Substrates for double-end binding experiments were generated by annealing complementary oligos to create 2-nt 5′ TA overhangs at both ends. The oligos were mixed in equimolar concentrations (10 mM) in STE buffer (100 mM NaCl, 10 mM Tris-HCl pH 8, 1 mM EDTA), heated, and slowly cooled. The substrates were then 5′-end labeled with [γ-^32^P]-ATP (Perkin Elmer) and T4 polynucleotide kinase and purified by native PAGE.

The sequence of the oligonucleotide substrate for DNA cleavage was designed with one of the preferred cutting sites from pUC19 (**Fig. 4a**,**b**) placed in the middle of a 44 bp duplex flanked by A4 ssDNA loops (**Fig. 4f**). The 96-nt self-complementary oligonucleotide was first purified using a 15% polyacrylamide-urea gel (Invitrogen) and then labeled with [γ-^32^P]-ATP (Perkin Elmer) and T4 polynucleotide kinase. The purified oligonucleotide was then annealed by heating and slow cooling. T4 DNA ligase was then used to seal the DNA nick, and the circular ligated oligonucleotide was purified from linear unligated molecules using another 15% polyacrylamide-urea gel. The circular oligonucleotide was then reannealed by heating and slow cooling.

### Electrophoretic mobility shift assays

For hairpin substrates, binding reactions (10 μl) were carried out in 25 mM Tris-HCl pH 7.5, 7.5% glycerol, 100 mM NaCl, 2 mM DTT, 5 mM MgCl_2_, and 1 mg/ml BSA with 0.1 nM DNA and the indicated concentration of SPO11 complexes. Reactions were assembled on ice then incubated for 30 min at 30 °C and separated on a 6% DNA retardation gel (Invitrogen) at 100 V for 30 min. For binding to the oligonucleotide cleavage substrate, SPO11 complex was incubated with 0.1 nM DNA at 4 °C for 30 min in a buffer containing 25 mM HEPES-NaOH pH 7.5, 50 mM NaCl, 5 mM MgCl_2_, 7.5% glycerol, 1 mg/ml BSA, and 2 mM DTT. A 6% DNA retardation gel was pre-run at 100 V for 20 min with 0.5′ Tris-borate supplemented with 5 mM MgCl_2_ as the running buffer, then the binding reactions were loaded and separated at 100 V for 40 min. For both substrates, gels were dried, exposed to autoradiography plates, and visualized by phosphor imaging. Quantification was performed using GelBandFitter^57^, and apparent *K*_*d*_ values were calculated by nonlinear regression in GraphPad Prism 10.

### Atomic force microscopy imaging

Linear plasmids for AFM were prepared by treatment of pUC19 with NdeI. SPO11–TOP6BL complexes were diluted to a final concentration of 4 nM in the presence of 1 nM (molecules) DNA in 25 mM HEPES-NaOH pH 6.8, 5 mM MgCl_2_, 50 mM NaCl, 10% glycerol. Mixtures were incubated at 30 °C for 30 min. A volume of 40 μl of the DNA-protein mixture was deposited onto pretreated mica with 3-amino-proply-trietoxy silane (APTES) (Pelco Mica Disc, V1, Ted Pella) for 5 min. The sample-deposited mica was rinsed with 1 ml molecular biology grade deionized water and dried gently with a stream of nitrogen. AFM images were captured using an Nanowizard V (JPK-Bruker) in AC Mode Imaging at room temperature. AFM probe (RFESPA-75, Bruker) with nominal frequencies of approximately 75 kHz and nominal spring constant of 3 N/m was used for imaging. Images were collected at a 2 Hz line rate with an image size of 2 × 2 μm at 512 × 512-pixel resolution. For data processing, images were exported with 3080-pixel resolution. The images were processed with JPK Data Processing software. ImageJ was used to quantify DNA bending angles.

### Mass photometry

Mass photometry experiments were conducted using a Refeyn On-eMP instrument (Oxford, UK). A ready-to-use 6-well sample cassettes (Refeyn) was placed at the center of a clean, ready-to-use sample carrier slide (Refeyn), with each well designated for a single measurement. To find the focus, 15 µl of fresh MP buffer (25 mM HEPES-NaOH pH 7.5, 150 mM NaCl, 2 mM DTT) was first loaded into the well. The focal position was identified and secured for the entire measurement using an autofocus system based on total internal reflection. Purified SPO11 or LZ-SPO11 complex was diluted to a concentration of 240 nM in MP buffer, either 2 µl or 1 µl of the diluted SPO11 or LZ-SPO11 complex was added to the buffer drop, resulting in a final protein concentration of 28 nM or 15 nM, respectively. After autofocus stabilization, movies were recorded for a duration of 60 s. Data acquisition was performed using AcquireMP (Refeyn Ltd) and data analysis was carried out with DiscoverMP (Refeyn Ltd). Contrast-to-mass calibration was performed using a BSA protein standard (Sigma), containing BSA monomers and dimers with molecular weights of 66.5 and 132 kDa. Statistical analysis was performed using DiscoverMP, where the distribution peaks were fitted with Gaussian functions to obtain the average molecular mass of each distribution component. Plotting was carried out using GraphPad Prism 10.

### DNA cleavage assays

Positively supercoiled pUC19 was prepared by incubating negatively supercoiled pUC19 (Thermo Scientific) with reverse gyrase (generous gift of S. Bahng in the K. Marians laboratory, MSK), followed by purification using a QIAquick PCR purification kit (QIAGEN) and verification via chloroquine-containing agarose gel electrophoresis as described^58^. Relaxed covalently closed pUC19 was generated by incubating negatively supercoiled pUC19 with *E. coli* Topoisomerase I (NEB) and purified using a QIAquick PCR purification kit (QIAGEN). Linearized substrates were prepared by double digestion of pUC19 with EcoRI and SspI, followed by agarose gel extraction using a QIAquick Gel Extraction Kit (QIAGEN). For quantification, the linearized substrates were labeled at both 5′ ends using [γ-^32^P]-ATP (Perkin Elmer) and T4 polynucleotide kinase (NEB).

Typical plasmid reactions (70 μl) were carried out in a buffer containing 25 mM HEPES-NaOH pH 7.5, 1 mM DTT, 0.1 mg/ml BSA (Sigma), 1 mM MnCl_2_, and 4 ng/µl pUC19 DNA. The reaction buffer was prepared on ice, and the recombinant proteins (usually 75–100 nM) were then added on ice. Unless otherwise stated, the reactions were incubated at 37 °C for the specified times. At each indicated time point, 11 µl of the reaction mixture was removed and terminated with 3 μl of STOP solution (375 mM EDTA, 5% SDS) and 1 μl of proteinase K (Thermo Scientific, 20 mg/ml), followed by incubation for 30 min at 50 °C.

The reaction products were then mixed with 6× DNA loading dye (NEB) and separated by electrophoresis on a 1.2% agarose gel (Lonza) in TAE buffer. Using a customized vertical agarose gel system (gel length 14.5 cm, CBS Scientific), separation was carried out for 90 min at 80 V. The gels were subsequently stained with SYBR Gold (Invitrogen) and imaged using a ChemiDoc MP imaging system (Bio-Rad).

Gel images were quantified using GelBandFitter. Cleavage activity is expressed as percent of substrate utilized, i.e., the fraction of DNA molecules that had sustained at least one detectable nick or DSB. It is important to note that this underestimates total strand cleavage when DNA molecules sustain multiple breaks. Substrate utilization was calculated as the percentage of nicks and DSBs relative to the total DNA in each lane, subtracting any nicked DNA present in control (no protein) reactions as background. When multiple DSBs were clearly present, i.e., where clear smearing appeared on the gel at later time points, we quantified substrate utilization for those lanes instead as the disappearance of the supercoiled band compared to time zero. For plotting purposes, substrate utilization time courses were fitted to two-phase association curves by nonlinear regression in Prism 10. These curves are only intended to aid in visualization of trends, and should not be viewed as a theoretically valid way to estimate underlying rate parameters.

For oligonucleotide cleavage assays, reactions were performed in a buffer containing 25 mM HEPES-NaOH pH 7.5), 5 mM MnCl_2_ (or 5 mM MgCl_2_), 0.1 mg/ml BSA, 1 mM DTT, and 0.02% NP40. Mixtures containing 10 nM SPO11 complexes and 0.5 nM radiolabeled oligonucleotide were assembled at 4 °C and then immediately transferred to 37 °C to initiate the reaction. At each specified time point, 5 µl of the reaction mixture was taken out and rapidly quenched with 1.2 µl of 375 mM EDTA. The mixture was then digested with proteinase K for 30 minutes at 50 °C. Samples were loaded onto a 15% polyacrylamide-urea gel in 1× TBE running buffer and electrophoresed at 180 V for 50 min. Gels were then dried, exposed to autoradiography plates, visualized by phosphor imaging, and analyzed using GelBandFitter. For experiments mixing SPO11-Y138F and SPO11-E224A, 5 nM of each mutant complex was used. For SPO11-Y138F alone, 10 nM of the mutant complex were used.

### Assays for covalent protein-DNA complexes

For the CsCl ultracentrifugation and immunoprecipitation assays, cleavage reactions (60 µl) were carried out in a buffer containing 25 mM HEPES-NaOH pH 7.5, 1 mM DTT, 0.1 mg/ml BSA, 5 mM MnCl_2_, and 4 ng/µl pUC19 DNA. The reaction buffer was prepared on ice, and the recombinant proteins (325 nM final concentration) were then added on ice. The reactions were incubated at 37 °C for 11 min. To terminate the reaction for the ultracentrifugation assay, 50 µl of the reaction mixture was combined with 83.3 µl of a stop solution to yield final concentrations of 30 mM EDTA, 0.5% Sarkosyl, 5 M guanidine HCl. To terminate the reaction for the immunoprecipitation assay, 50 µl of the reaction mixture was combined with 12.5 µl of a stop solution to yield final concentrations of 37.5 mM EDTA, 0.5% SDS.

An adaptation of the ICE (in vivo complex of enzymes) assay was used for the immunodetection of proteins covalently bound to DNA^59^. In a new 5 ml centrifuge tube, 2 ml of 150% (w/v) CsCl solution was added, then 2 ml of buffer containing 10 mM Tris-HCl pH 8.0, 0.1 mM EDTA, and 0.5% Sarkosyl was layered on top. The stopped reaction mixture (133.3 µl) was then layered on top, and buffer was added until the tubes were full. The tubes were sealed, placed in a TN-1865 ultracentrifuge rotor (Thermo Scientific), and centrifuged at 42,000 rpm (∼157,000 g) for 17.5 h at 24 °C in a Sorvall wX+ Ultra Series ultracentrifuge (Thermo Scientific). The resulting DNA pellets with covalently bound proteins were washed with 70% ethanol and dissolved in 1× TE buffer (10 mM Tris-HCl pH 8.0, 0.1 mM EDTA) for 2 h at room temperature. Each sample was mixed with one-third volume of 25 mM sodium phosphate pH 6.5 buffer, then applied to a 0.45 µm nitrocellulose membrane (Bio-Rad) using a slot-blot vacuum manifold (Bio-Rad). Recombinant SPO11 complex (∼10 ng total protein) was applied to adjacent slots as a control for detection. SPO11 protein was immunodetected using an anti-Flag-HRP monoclonal antibody (mouse, 1:1000, Sigma), followed by incubation with ECL Prime western blotting detection reagent (Amersham) and detection using a ChemiDoc MP imaging system (Bio-Rad).

For immunoprecipitation, the stopped reaction mixture was mixed with 237.5 µl of binding buffer (25 mM HEPES-NaOH pH 7.5, 500 mM NaCl, 5 mM EDTA, 1% Triton X-100) and 50 µl of anti-Flag magnetic agarose (Pierce), which had been pre-washed twice with binding buffer. The mixture was incubated at room temperature with gentle shaking at 1000 rpm for 60 min. After incubation, the magnetic agarose was washed twice with wash buffer (25 mM HEPES-NaOH pH 7.5, 150 mM NaCl, 5 mM EDTA, 1% Triton X-100), then 30 µl of 2× Laemmli sample buffer (BioRad) and 1 µl of proteinase K (Thermo Scientific, 20 mg/ml) were added, followed by incubation for 30 min at 50 °C. The products were purified using the QIAquick PCR purification kit (QIAGEN), mixed with 6× DNA loading dye (NEB), and separated by agarose gel electrophoresis as in the DNA cleavage assays. The gels were subsequently stained with SYBR Gold (Invitrogen) and imaged using a ChemiDoc MP imaging system (Bio-Rad).

To investigate 5′ or 3′ SPO11 attachment, 10 µg of pUC19 was cleaved with EcoRI and labeled at the 5′ ends using T4 polynucleotide kinase (NEB) with [γ-^32^P]-ATP or at the 3′ ends using Klenow (NEB) with [a-^32^P]-dATP and [a-^32^P]-TTP. Following labeling, the substrates were further digested with SspI to produce single-end labeled fragments, which were subsequently purified by agarose gel electrophoresis and extraction using QIAquick Gel Extraction Kit (QIAGEN). Cleavage reactions were performed at a substrate concentration of 2.3 nM (0.87 ng/µl) in the presence of 25 mM HEPES-NaOH pH 7.5, 1 mM DTT, 0.1 mg/ ml BSA, and 5 mM MnCl_2_. The reaction buffer was prepared on ice, and recombinant proteins (325 nM) were added on ice. Reactions were incubated at 37 °C for 2 h. To terminate the reactions, 70 µl of the reaction mixture was combined with 17.5 µl of stop solution to achieve final concentrations of 37.5 mM EDTA and 0.5% SDS. A 43.75 µl aliquot (half) of the stopped reaction mixture was treated with 1 µl proteinase K and incubated at 50 °C for 1 h. Samples, with or without proteinase K treatment, were then mixed with an equal volume of 2× Urea TBE loading buffer (Invitrogen) and heated at 85 °C for 5 min. Subsequently, 10 µl of each sample was loaded onto a prewarmed 6% TBE-urea gel (Invitrogen) at 60 °C and electrophoresed at 180 V for 30 min. Gels were dried, exposed to autoradiography plates, and visualized using phosphor imaging.

### TDP2-seq

In vitro cleavage assays for sequencing were conducted in 70 µl reaction buffer containing 25 mM HEPES-NaOH pH 7.5, 1 mM DTT, 0.1 mg/ml BSA and 1 mM MnCl_2_. For cleavage of pUC19, reaction buffer was mixed with 4 ng/µl DNA (final concentration) on ice, then recombinant SPO11 (100 nM) was added and the mixture was incubated at 37 °C for 30 min followed by inactivation with 1 µl proteinase K solution (Thermo Scientific, 20 mg/ml) and 30 min incubation at 50 °C. Bacterial and yeast genomic DNA was purified by extraction with phenol-chloroform-isoamyl alcohol (25:24:1; Thermo Fisher) and ethanol precipitation from *E. coli* DH5α cells (Takara) or exponentially growing *S. cerevisiae* cells of the S288C strain background. For genomic DNA cleavage, reaction buffer was mixed with 2.5 µg DNA on ice. Recombinant SPO11 (300 nM) was then added and the mixture was incubated at 37 °C for 60 min and inactivated as above.

Sequencing libraries were prepared using a modified version of the S1-sequencing protocol^60-63^. In brief, mouse testes were dissociated, and cells were embedded in agarose plugs as described^63^. For mapping in vitro SPO11 cleavage sites, DNA digested by SPO11 in vitro was embedded in plugs together with wild type C57BL/6J mouse testis cells (0.5–1 million cells per plug), whose DNA acted as carrier during library preparation. Two plugs were prepared for each experiment. Following proteinase K and RNase A treatment, plugs were equilibrated in TDP2 buffer according to the TopoGEN protocol (50 mM Tris-HCl pH 8, 0.15 M NaCl, 10 mM MgCl_2_, 0.5 mM DTT, 30 μg/ml BSA, 2 mM ATP) and incubated with purified human TDP2 protein (490 pmol per plug, TopoGEN) at 37 °C for 30 min for removal of covalently linked SPO11 from DNA ends. Subsequent steps (end fill-in with T4 DNA polymerase (NEB), ligation to biotinylated adaptors, DNA purification, DNA shearing, ligation to a second adaptor at the sheared end, and PCR amplification) were all performed as described^63^ with the following minor modifications: 1) to reduce the loss of very small DNA fragments from diffusion out of the plugs, the washing step after ligation of the first adaptor was reduced from overnight to 1 h at 4 °C for experiments with pUC19; 2) the NEBNext End Repair Module (NEB, E6050S) was used for repair of DNA ends after shearing. DNA libraries were sequenced on the Illumina NextSeq 1000 platform (paired-end, 50 bp) at the Integrated Genomics Operation at Memorial Sloan Kettering Cancer Center.

Sequencing reads were mapped against the pUC19, *E. coli* (ASM584v2), *S. cerevisiae* (sacCer3) or mouse (mm10) reference sequence using modified versions of previously published custom pipelines^62^. Briefly, reads were mapped using bowtie2^64^ with parameters -N 1 -X 1000. Uniquely mapped reads were extracted and assigned to the nucleotide immediately next to the biotinylated adaptor. Statistical analyses were performed using R versions 4.2.3 and 4.3.2. Telomeres (500 bp at the ends of each chromosome) and rDNA (chrXII:459,400-460,900) in the *S. cerevisiae* genome were masked before downstream analyses. Mitochondrial DNA and the 2 µ plasmid were also excluded. Raw and processed TDP2-seq data were deposited at the Gene Expression Omnibus (GEO) (accession number GSE275291). Mapping statistics are found in **Supplemental Table 2**.

### Peak calling and base composition analyses

Top- and bottom-strand peaks were separately called as nucleotide positions with >3,000, >10 and >15 RPM of strand-specific read counts in pUC19, *E. coli*, and *S. cerevisiae* TDP2-seq data, respectively. Positions with less read counts than those located at –1 and +1 positions on top and bottom strands, respectively, were not defined as peaks because those reads could be false-positively enriched due to incomplete fill-in of 5′ overhangs at DNA ends before adaptor ligation. Base compositions were calculated for the strand from which TDP2-seq read was sequenced, i.e., bottom strand peaks were analyzed with reorientation so that the nucleotide immediately next to the biotinylated adaptor should be located at –1 position relative to dyad axis on top strand. Sequence logos were generated using ggseqlogo^65^, with base composition biases corrected for the genome-average G+C content. For *S. cerevisiae* Spo11 in vivo data, the local G+C content around mapped DSB sites (47.8%) was used instead because it differed substantially from the genome average (38.5%). Motif matches in **Fig. 4e** were identified using dreg from the European Molecular Biology Open Software Suite^66^ with default parameters.

### AlphaFold3

The pre-DSB SPO11 dimer models were generated using the Alphafold3 online service (https://golgi.sandbox.google.com)^7^. The input comprised two full-length mouse SPO11 (Uniprot: Q9WTK8-1) and TOP6BL (Uniprot: J3QMY9-1) proteins, two complementary DNA sequences (**Supplemental Table S1**), and four magnesium ions. Multiple DNA sequences were used to evaluate dependence of the model on the DNA composition. The sequences were selected to represent different SPO11 cleavage sites in yeast genomic DNA, and they varied in length from 36 to 44 bp. Each session produced five top-ranked models. We selected one representative model for figure preparation; this model’s coordinates are provided in **Supplemental File 1**. Structural alignments, analysis, and figure generation were performed in Chimera^67^ and ChimeraX^68^.

## Data availability

Raw and processed TDP2-seq data are available at GEO under accession number GSE275291. The AlphaFold3 model used to generate most of the figures is provided in .pdb format as Supplemental Data.

## Acknowledgments

This article is subject to the Open Access to Publications policy of the Howard Hughes Medical Institute (HHMI). HHMI lab heads have previously granted a nonexclusive CC BY 4.0 license to the public and a sublicensable license to HHMI in their research articles. Pursuant to those licenses, the author-accepted manuscript of this article can be made freely available under a CC BY 4.0 license immediately upon publication. We thank C. Claeys Bouuaert and M. Tong for discussion and sharing of unpublished data. We thank K. Marians, S. Bahng, C. Lee, D. Remus, and members of the Keeney and D. Patel labs for discussions and experimental advice. We thank M. Lu (Keeney lab) for providing a *Mre11* conditional knockout mouse. We thank the MSK Integrated Genomics Operation for sequencing and Y.H. Kim in the Molecular Cytology Core Facility for AFM imaging. MSK core facilities are supported by National Cancer Institute cancer center support grant P30 CA08748. ZZ is supported in part by a Bruce Charles Forbes Predoctoral Fellowship. KL was supported in part by the Damon Runyon Cancer Research Foundation (DRG-[2389-20] to KL). MA was supported in part by an EMBO Long Term Fellowship (ALTF 905-2019 to MA). This work was supported by NIH grant R01 HD110120 (to S. Keeney and D. Patel); an MSK Basic Research Innovation Award (BRIA, to SK and D. Patel); and NIH grant R35 GM118092 (to S. Keeney). S. Keeney is an HHMI investigator. The funders had no role in study design, data collection and analysis, decision to publish or preparation of the manuscript.

## Author contributions

S. Keeney conceived the project; ZZ, LZ, MA, KL, DO and S. Keeney designed experiments; ZZ, LZ, MA, and DO performed experiments; ZZ, LZ, MA, KL, SY, DO and S. Keeney analyzed data; S. Kim contributed unpublished reagents and data; S. Keeney supervised the research; ZZ, KL, MA, and S. Keeney secured funding; S. Keeney wrote the paper; all authors edited the manuscript.

## References

1 Lam, I. & Keeney, S. Mechanism and regulation of meiotic recombination initiation. Cold Spring Harb Perspect Biol 7, a016634 (2014). 10.1101/cshperspect.a016634

2 Hinch, R., Donnelly, P. & Hinch, A. G. Meiotic DNA breaks drive multifaceted mutagenesis in the human germ line. Science 382, eadh2531 (2023). 10.1126/science.adh2531

3 Handel, M. A. & Schimenti, J. C. Genetics of mammalian meiosis: regulation, dynamics and impact on fertility. Nat Rev Genet 11, 124–136 (2010). 10.1038/nrg2723

4 Keeney, S., Lange, J. & Mohibullah, N. Self-organization of meiotic recombination initiation: general principles and molecular pathways. Annu Rev Genet 48, 187–214 (2014). 10.1146/annurev-genet-120213-092304

5 Yu, Y. et al. Cryo-EM structure of the Spo11 core complex bound to DNA. bioRxiv (2023). 10.1101/2023.10.31.564985

6 Claeys Bouuaert, C. et al. Structural and functional characterization of the Spo11 core complex. Nat Struct Mol Biol 28, 92–102 (2021). 10.1038/s41594-020-00534-w

7 Abramson, J. et al. Accurate structure prediction of biomolecular interactions with AlphaFold 3. Nature 630, 493–500 (2024). 10.1038/s41586-024-07487-w

8 Hunter, N. Meiotic recombination: The essence of heredity. Cold Spring Harb Perspect Biol 7, 361–395 (2015). 10.1101/cshperspect.a016618

9 Zickler, D. & Kleckner, N. Meiosis: Dances between homologs. Annu Rev Genet 57, 1–63 (2023). 10.1146/annurev-genet-061323-044915

10 Jiao, Y. et al. A TOP6BL mutation abolishes meiotic DNA double-strand break formation and causes human infertility. Sci Bull (Beijing) 65, 2120–2129 (2020). 10.1016/j.scib.2020.08.026

11 Robert, T. et al. The TopoVIB-Like protein family is required for meiotic DNA double-strand break formation. Science 351, 943–949 (2016). 10.1126/science.aad5309

12 Baudat, F., Manova, K., Yuen, J. P., Jasin, M. & Keeney, S. Chromosome synapsis defects and sexually dimorphic meiotic progression in mice lacking Spo11. Mol Cell 6, 989–998 (2000). 10.1016/s1097-2765(00)00098-8

13 Romanienko, P. J. & Camerini-Otero, R. D. The mouse Spo11 gene is required for meiotic chromosome synapsis. Molecular Cell 6, 975–987 (2000). 10.1016/S1097-2765(00)00097-6

14 Bergerat, A. et al. An atypical topoisomerase II from Archaea with implications for meiotic recombination. Nature 386, 414–417 (1997). 10.1038/386414a0

15 Keeney, S., Giroux, C. N. & Kleckner, N. Meiosis-specific DNA double-strand breaks are catalyzed by Spo11, a member of a widely conserved protein family. Cell 88, 375–384 (1997). 10.1016/s0092-8674(00)81876-0

16 Corbett, K. D., Benedetti, P. & Berger, J. M. Holoenzyme assembly and ATP-mediated conformational dynamics of topoisomerase VI. Nat Struct Mol Biol 14, 611–619 (2007). 10.1038/nsmb1264

17 Graille, M. et al. Crystal structure of an intact type II DNA topoisomerase: insights into DNA transfer mechanisms. Structure 16, 360–370 (2008). 10.1016/j.str.2007.12.020

18 Buhler, C., Lebbink, J. H., Bocs, C., Ladenstein, R. & Forterre, P. DNA topoisomerase VI generates ATP-dependent double-strand breaks with two-nucleotide overhangs. J Biol Chem 276, 37215–37222 (2001). 10.1074/jbc.M101823200

19 Nichols, M. D., DeAngelis, K., Keck, J. L. & Berger, J. M. Structure and function of an archaeal topoisomerase VI subunit with homology to the meiotic recombination factor Spo11. Embo J 18, 6177–6188 (1999). 10.1093/emboj/18.21.6177

20 Neale, M. J., Pan, J. & Keeney, S. Endonucleolytic processing of covalent protein-linked DNA double-strand breaks. Nature 436, 1053–1057 (2005). 10.1038/nature03872

21 Vrielynck, N. et al. A DNA topoisomerase VI-like complex initiates meiotic recombination. Science 351, 939–943 (2016). 10.1126/science.aad5196

22 Malone, R. E. et al. Isolation of mutants defective in early steps of meiotic recombination in the yeast Saccharomyces cerevisiae. Genetics 128, 79–88 (1991). 10.1093/genetics/128.1.79

23 Kee, K. & Keeney, S. Functional interactions between SPO11 and REC102 during initiation of meiotic recombination in Saccharomyces cerevisiae. Genetics 160, 111–122 (2002). 10.1093/genetics/160.1.111

24 Brinkmeier, J., Coelho, S., de Massy, B. & Bourbon, H. M. Evolution and diversity of the TopoVI and TopoVI-like subunits with extensive divergence of the TOPOVIBL subunit. Mol Biol Evol 39 (2022). 10.1093/molbev/msac227

25 Yeh, H. Y., Lin, S. W., Wu, Y. C., Chan, N. L. & Chi, P. Functional characterization of the meiosis-specific DNA double-strand break inducing factor SPO-11 from C. elegans. Scientific reports 7, 2370 (2017). 10.1038/s41598-017-02641-z

26 An, X. J., Deng, Z. Y. & Wang, T. OsSpo11-4, a rice homologue of the archaeal TopVIA protein, mediates double-strand DNA cleavage and interacts with OsTopVIB. PLoS One 6, e20327 (2011). 10.1371/journal.pone.0020327

27 Shingu, Y., Mikawa, T., Onuma, M., Hirayama, T. & Shibata, T. A DNA-binding surface of SPO11-1, an Arabidopsis SPO11 orthologue required for normal meiosis. FEBS J 277, 2360–2374 (2010). 10.1111/j.1742-4658.2010.07651.x

28 Wu, H., Gao, J., Sharif, W. D., Davidson, M. K. & Wahls, W. P. Purification, folding, and characterization of Rec12 (Spo11) meiotic recombinase of fission yeast. Protein Expr Purif 38, 136–144 (2004). 10.1016/j.pep.2004.07.012

29 Arter, M. & Keeney, S. Divergence and conservation of the meiotic recombination machinery. Nat Rev Genet 25, 309–325 (2024). 10.1038/s41576-023-00669-8

30 Jolivet, S., Vezon, D., Froger, N. & Mercier, R. Non conservation of the meiotic function of the Ski8/Rec103 homolog in Arabidopsis. Genes to Cells 11, 615–622 (2006). 10.1111/j.1365-2443.2006.00972.x

31 Diagouraga, B. et al. The TOPOVIBL meiotic DSB formation protein: new insights from its biochemical and structural characterization. Nucleic Acids Res (2024). 10.1093/nar/gkae587

32 Johnson, D. et al. Concerted cutting by Spo11 illuminates meiotic DNA break mechanics. Nature 594, 572–576 (2021). 10.1038/s41586-021-03389-3

33 de Massy, B., Rocco, V. & Nicolas, A. The nucleotide mapping of DNA double-strand breaks at the CYS3 initiation site of meiotic recombination in Saccharomyces cerevisiae. EMBO J. 14, 4589–4598 (1995). 10.1002/j.1460-2075.1995.tb00138.x

34 Keeney, S. & Kleckner, N. Covalent protein-DNA complexes at the 5’ strand termini of meiosis-specific double-strand breaks in yeast. Proc Natl Acad Sci USA 92, 11274–11278 (1995). 10.1073/pnas.92.24.11274

35 Liu, J., Wu, T. C. & Lichten, M. The location and structure of double-strand DNA breaks induced during yeast meiosis: evidence for a covalently linked DNA-protein intermediate. EMBO J 14, 4599–4608 (1995). 10.1002/j.1460-2075.1995.tb00139.x

36 Buhler, C., Gadelle, D., Forterre, P., Wang, J. C. & Bergerat, A. Reconstitution of DNA topoisomerase VI of the thermophilic archaeon Sulfolobus shibatae from subunits separately overexpressed in Escherichia coli. Nucleic Acids Res 26, 5157–5162 (1998). 10.1093/nar/26.22.5157

37 Corbett, K. D. & Berger, J. M. Structural dissection of ATP turnover in the prototypical GHL ATPase TopoVI. Structure 13, 873–882 (2005). 10.1016/j.str.2005.03.013

38 Wendorff, T. J. & Berger, J. M. Topoisomerase VI senses and exploits both DNA crossings and bends to facilitate strand passage. eLife 7 (2018). 10.7554/eLife.31724

39 Dutta, R. & Inouye, M. GHKL, an emergent ATPase/kinase superfamily. Trends Biochem Sci 25, 24–28 (2000). 10.1016/s0968-0004(99)01503-0

40 Pan, J. et al. A hierarchical combination of factors shapes the genome-wide topography of yeast meiotic recombination initiation. Cell 144, 719–731 (2011). 10.1016/j.cell.2011.02.009

41 Murakami, H. & Nicolas, A. Locally, meiotic double-strand breaks targeted by Gal4BD-Spo11 occur at discrete sites with a sequence preference. Mol Cell Biol 29, 3500–3516 (2009). 10.1128/MCB.00088-09

42 Gittens, W. H. et al. A nucleotide resolution map of Top2-linked DNA breaks in the yeast and human genome. Nature communications 10, 4846 (2019). 10.1038/s41467-019-12802-5

43 Kim, S. et al. The MRE11-RAD50-NBS1 complex both starts and extends DNA end resection in mouse meiosis. bioRxiv, 2024.2008.2017.608390 (2024). 10.1101/2024.08.17.608390

44 Prieler, S. et al. Spo11 generates gaps through concerted cuts at sites of topological stress. Nature 594, 577–582 (2021). 10.1038/s41586-021-03632-x

45 O’Shea, E. K., Klemm, J. D., Kim, P. S. & Alber, T. X-ray structure of the GCN4 leucine zipper, a two-stranded, parallel coiled coil. Science 254, 539–544 (1991). 10.1126/science.1948029

46 Claeys Bouuaert, C. et al. DNA-driven condensation assembles the meiotic DNA break machinery. Nature 592, 144–149 (2021). 10.1038/s41586-021-03374-w

47 Lange, J. et al. The landscape of mouse meiotic double-strand break formation, processing, and repair. Cell 167, 695–708 e616 (2016). 10.1016/j.cell.2016.09.035

48 Tischfield, S. E. & Keeney, S. Scale matters: the spatial correlation of yeast meiotic DNA breaks with histone H3 trimethylation is driven largely by independent colocalization at promoters. Cell cycle 11, 1496–1503 (2012). 10.4161/cc.19733

49 Murakami, H. et al. Multilayered mechanisms ensure that short chromosomes recombine in meiosis. Nature 582, 124–128 (2020). 10.1038/s41586-020-2248-2

50 Brick, K., Smagulova, F., Khil, P., Camerini-Otero, R. D. & Petukhova, G. V. Genetic recombination is directed away from functional genomic elements in mice. Nature 485, 642–645 (2012). 10.1038/nature11089

51 Pratto, F. et al. Meiotic recombination mirrors patterns of germline replication in mice and humans. Cell 184, 4251–4267 e4220 (2021). 10.1016/j.cell.2021.06.025

52 Buis, J. et al. Mre11 nuclease activity has essential roles in DNA repair and genomic stability distinct from ATM activation. Cell 135, 85–96 (2008). 10.1016/j.cell.2008.08.015

53 Schonhoff, S. E., Giel-Moloney, M. & Leiter, A. B. Neurogenin 3-expressing progenitor cells in the gastrointestinal tract differentiate into both endocrine and non-endocrine cell types. Dev Biol 270, 443–454 (2004). 10.1016/j.ydbio.2004.03.013

54 Bellani, M. A., Boateng, K. A., McLeod, D. & Camerini-Otero, R. D. The expression profile of the major mouse SPO11 isoforms indicates that SPO-11beta introduces double strand breaks and suggests that SPO11alpha has an additional role in prophase in both spermatocytes and oocytes. Mol Cell Biol 30, 4391–4403 (2010). 10.1128/MCB.00002-10

55 Kauppi, L. et al. Distinct properties of the XY pseudoautosomal region crucial for male meiosis. Science 331, 916–920 (2011). 10.1126/science.1195774

56 Giannattasio, T. et al. The proper interplay between the expression of Spo11 splice isoforms and the structure of the pseudoautosomal region promotes XY chromosomes recombination. Cell Mol Life Sci 80, 279 (2023). 10.1007/s00018-023-04912-7

57 Mitov, M. I., Greaser, M. L. & Campbell, K. S. GelBandFitter--a computer program for analysis of closely spaced electrophoretic and immunoblotted bands. Electrophoresis 30, 848–851 (2009). 10.1002/elps.200800583

58 Hayama, R., Bahng, S., Karasu, M. E. & Marians, K. J. The MukB-ParC interaction affects the intramolecular, not intermolecular, activities of topoisomerase IV. J Biol Chem 288, 7653–7661 (2013). 10.1074/jbc.M112.418087

59 Anand, J., Sun, Y., Zhao, Y., Nitiss, K. C. & Nitiss, J. L. Detection of Topoisomerase Covalent Complexes in Eukaryotic Cells. Methods Mol Biol 1703, 283–299 (2018). 10.1007/978-1-4939-7459-7_20

60 Mimitou, E. P. & Keeney, S. S1-seq Assay for Mapping Processed DNA Ends. Methods Enzymol 601, 309–330 (2018). 10.1016/bs.mie.2017.11.031

61 Mimitou, E. P., Yamada, S. & Keeney, S. A global view of meiotic double-strand break end resection. Science 355, 40–45 (2017). 10.1126/science.aak9704

62 Yamada, S. et al. Molecular structures and mechanisms of DNA break processing in mouse meiosis. Genes Dev 34, 806–818 (2020). 10.1101/gad.336032.119

63 Kim, S., Yamada, S., Maekawa, K. & Keeney, S. Optimized methods for mapping DNA double-strand-break ends and resection tracts and application to meiotic recombination in mouse spermatocytes. bioRxiv (2024). 10.1101/2024.08.10.606181

64 Langmead, B., Trapnell, C., Pop, M. & Salzberg, S. L. Ultrafast and memory-efficient alignment of short DNA sequences to the human genome. Genome Biol 10, R25 (2009). 10.1186/gb-2009-10-3-r25

65 Wagih, O. ggseqlogo: a versatile R package for drawing sequence logos. Bioinformatics 33, 3645–3647 (2017). 10.1093/bioinformatics/btx469

66 Rice, P., Longden, I. & Bleasby, A. EMBOSS: the European Molecular Biology Open Software Suite. Trends Genet 16, 276–277 (2000). 10.1016/s0168-9525(00)02024-2

67 Pettersen, E. F. et al. UCSF Chimera--a visualization system for exploratory research and analysis. J Comput Chem 25, 1605–1612 (2004). 10.1002/jcc.20084

68 Pettersen, E. F. et al. UCSF ChimeraX: Structure visualization for researchers, educators, and developers. Protein Sci 30, 70–82 (2021). 10.1002/pro.3943

69 Nore, A. et al. TOPOVIBL-REC114 interaction regulates meiotic DNA double-strand breaks. Nature communications 13, 7048 (2022). 10.1038/s41467-022-34799-0

70 Li, J., Chiu, T. P. & Rohs, R. Predicting DNA structure using a deep learning method. Nature communications 15, 1243 (2024). 10.1038/s41467-024-45191-5

71 Li, J. & Rohs, R. Deep DNAshape webserver: prediction and real-time visualization of DNA shape considering extended k-mers. Nucleic Acids Res 52, W7–W12 (2024). 10.1093/nar/gkae433

72 Dereli, I. et al. Seeding the meiotic DNA break machinery and initiating recombination on chromosome axes. Nature communications 15, 2941 (2024). 10.1038/s41467-024-47020-1

73 Biot, M. et al. Principles of chromosome organization for meiotic recombination. Mol Cell 84, 1826–1841 e1825 (2024). 10.1016/j.molcel.2024.04.001

74 Yadav, V. K. & Claeys Bouuaert, C. Mechanism and control of meiotic DNA double-strand break formation in S. cerevisiae. Front Cell Dev Biol 9, 642737 (2021). 10.3389/fcell.2021.642737

75 Schmidt, B. H., Burgin, A. B., Deweese, J. E., Osheroff, N. & Berger, J. M. A novel and unified two-metal mechanism for DNA cleavage by type II and IA topoisomerases. Nature 465, 641–644 (2010). 10.1038/nature08974

76 Deweese, J. E., Burgin, A. B. & Osheroff, N. Human topoisomerase IIalpha uses a two-metal-ion mechanism for DNA cleavage. Nucleic Acids Res 36, 4883–4893 (2008). 10.1093/nar/gkn466

77 Pitts, S. L. et al. Use of divalent metal ions in the DNA cleavage reaction of topoisomerase IV. Nucleic Acids Res 39, 4808–4817 (2011). 10.1093/nar/gkr018

78 Diaz, R. L., Alcid, A. D., Berger, J. M. & Keeney, S. Identification of residues in yeast Spo11p critical for meiotic DNA double-strand break formation. Mol Cell Biol 22, 1106–1115 (2002). 10.1128/MCB.22.4.1106-1115.2002

