## Supplemental information for "Reconstitution of SPO11-dependent double-strand break formation"

**This pdf contains:**

Extended Data Figures 1–7
Supplemental Discussion
Supplemental Tables 1–2
Supplemental Figure 1

**Supplemental Data provided as a separate file:**

Supplemental File 1. AlphaFold3 model of SPO11–TOP6BL dimer bound to DNA

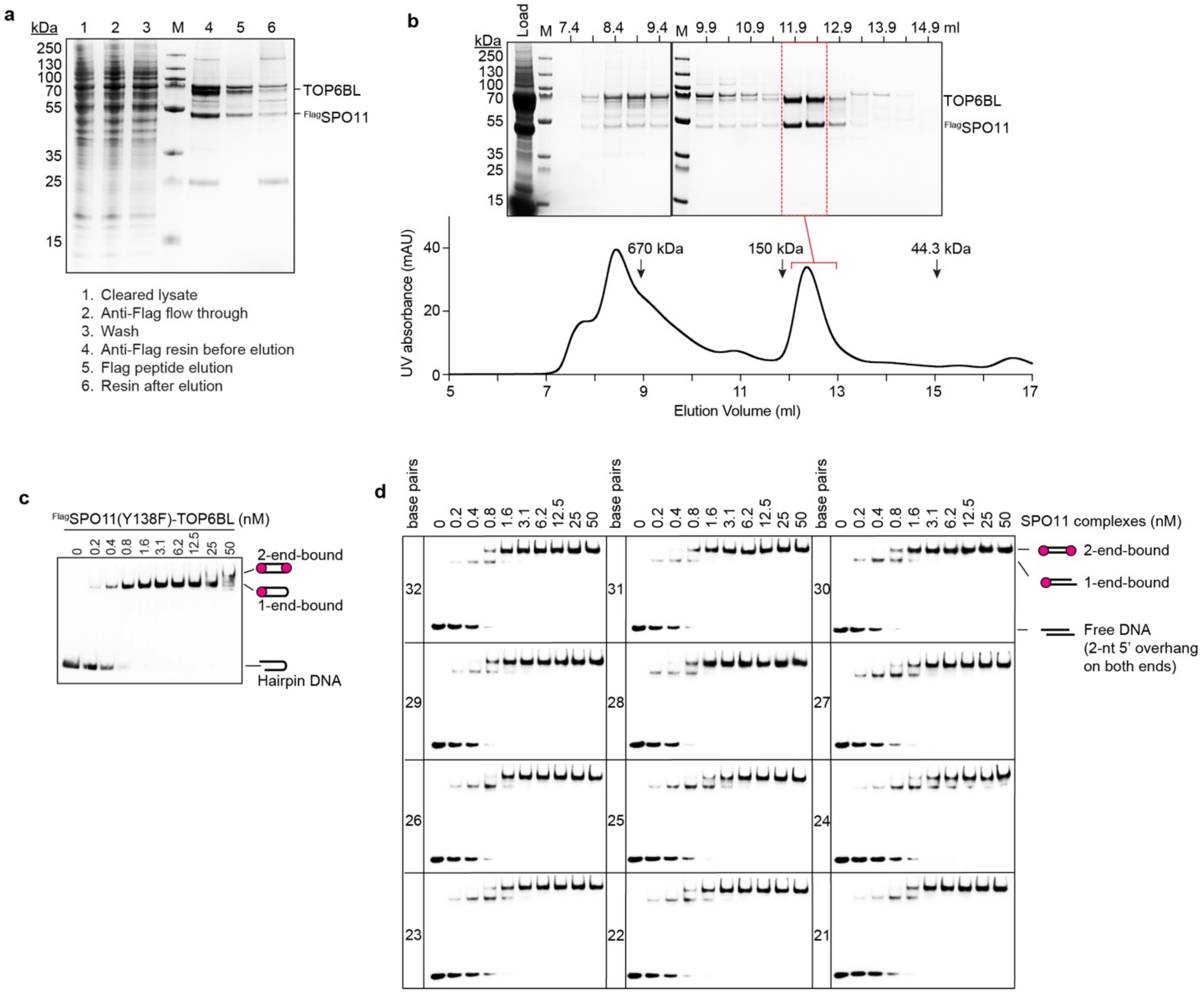


**Extended Data Fig. 1 | DNA binding by purified SPO11–TOP6BL complexes.**
**a**, Coomassie-stained SDS-PAGE gel of samples at the indicated steps from a representative purification.
**b**, Representative SEC profile. A Coomassie-stained SDS-PAGE gel of the indicated 0.5 ml fractions is above and UV profile is below. Elution positions of size standards analyzed in a separate calibration run are indicated (arrows). Pooled fractions are indicated in red.
**c**, Representative EMSA of SPO11-Y138F mutant complexes binding to a 25-bp hairpin substrate with a two-nucleotide 5′ overhang end. Quantification is in **Fig. 1d**.
**d**, DNA length dependence for double-end binding. Full EMSAs are shown for the experiment presented in **Fig. 1e**.

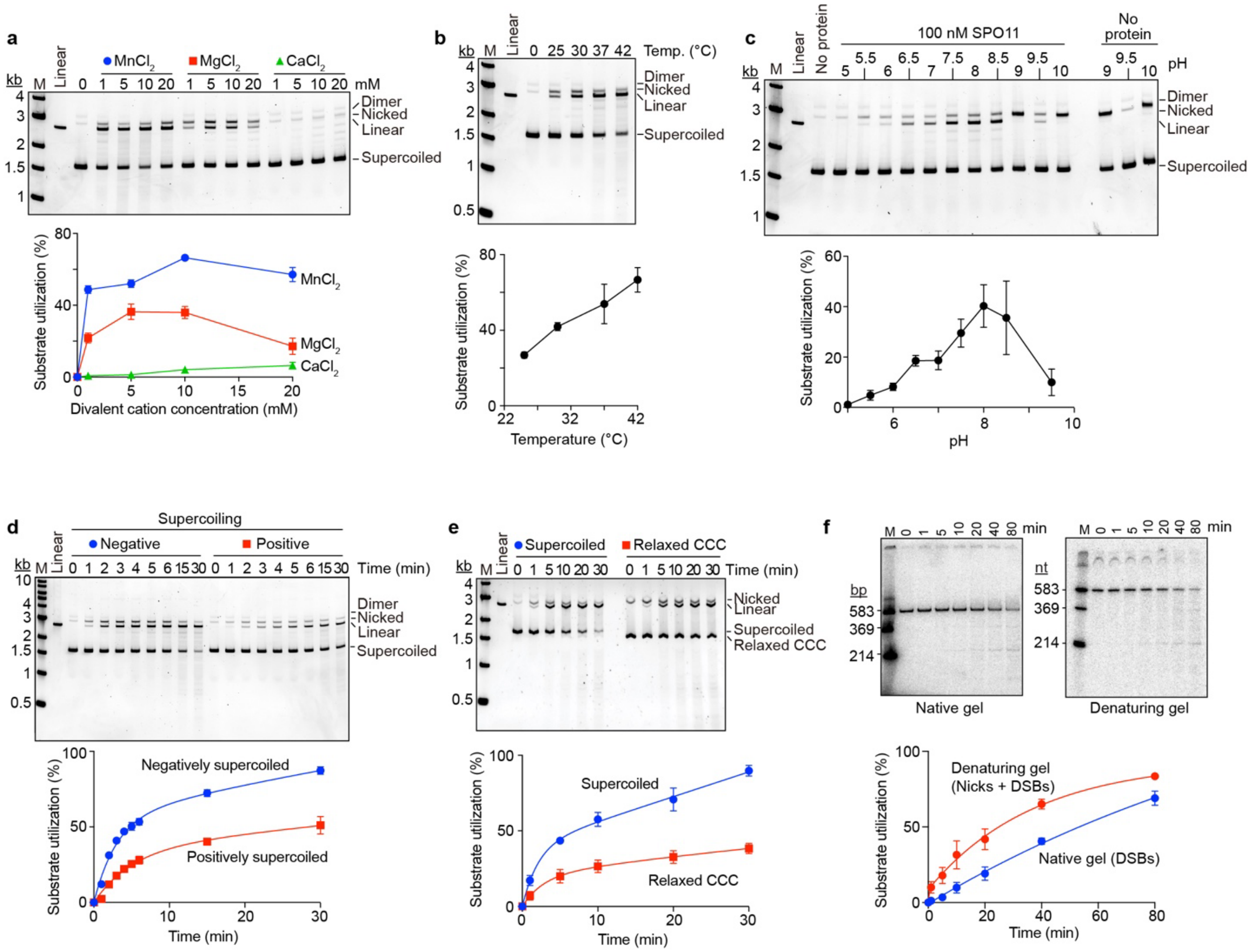

**Extended Data Fig. 2 | DNA cleavage optimization.**
In all panels, representative gels are above, quantification is below (mean ± s.d. of n=3 experiments).
**a**, Metal ion dependence of DNA cleavage. SPO11–TOP6BL complexes (100 nM) were incubated with 4 ng/μl pUC19 DNA in the presence of the indicated concentration of $MnCl_2$, $MgCl_2$, or $CaCl_2$.
**b**, Temperature dependence. Reactions contained 100 nM SPO11 complexes and 4 ng/μl pUC19 DNA with 1 mM $MnCl_2$.
**c**, pH dependence. Reactions contained 100 nM SPO11 complexes and 4 ng/μl pUC19 DNA with 1 mM $MnCl_2$. The pH 9.0 and pH 10.0 conditions resulted in a high background of SPO11-independent nicking (right lanes), so these samples were omitted from the quantification.
**d**, Comparison of positively and negatively supercoiled substrates. Reactions contained 100 nM SPO11 complexes and 4 ng/μl pUC19 DNA with 1 mM $MnCl_2$.
**e**, Comparison of relaxed covalently closed circle (CCC) and negatively supercoiled substrates. Reactions contained 100 nM SPO11 complexes and 4 ng/μl pUC19 DNA with 1 mM $MnCl_2$. For this experiment, reaction products were separated on agarose gels containing ethidium bromide.
**f**, Cleavage of a linear DNA substrate. Reactions contained 100 nM SPO11 complexes and 4 ng/μl of a linear DNA fragment from pUC19 (mix of cold and 5′ $^{32}P$-labeled on both ends) with 5 mM $MnCl_2$. Deproteinized reaction products were divided and aliquots were run separately on native PAGE to detect DSBs (left) and denaturing urea PAGE to detect both nicks and DSBs (right).

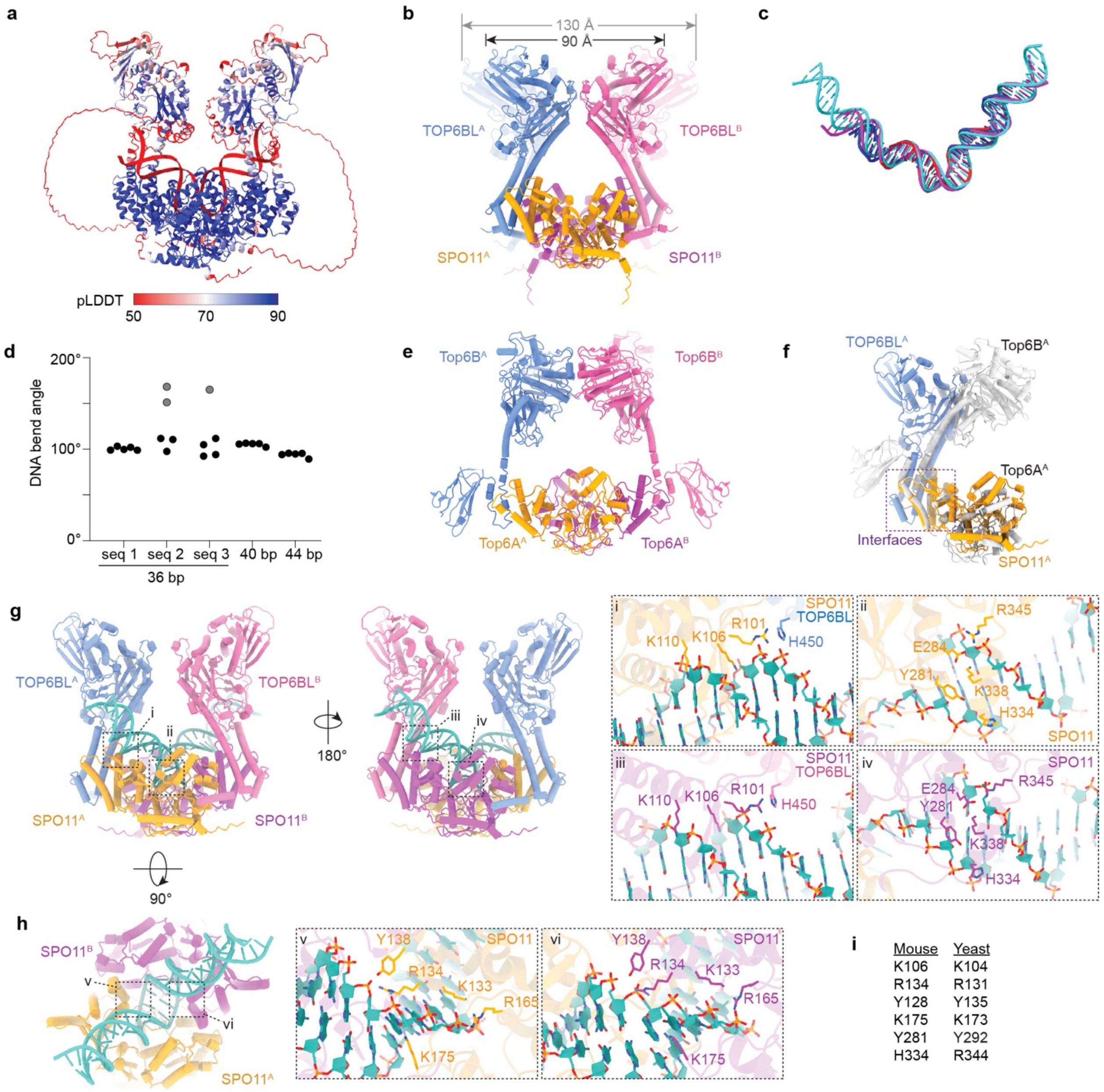


**Extended Data Fig. 3 | Computational models of the structure of dimeric SPO11–TOP6BL complexes bound to DNA.**

**a**, AlphaFold3 model colored according to pLDDT confidence score. Many of the models place a C-terminal helix from TOP6BL near the underside of the SPO11 from the other complex (i.e., TOP6BL$^{A}$ interacting with SPO11$^{B}$), preceded by a long linker predicted to be disordered. The C-terminal helix was previously shown to bind to REC114[69], but whether it can also interact with SPO11 as predicted by AlphaFold3 has not been verified.

**b**, Variation among AlphaFold3 models in distances between TOP6BL GHKL domains. The range in distances is shown for the models shown in **Supplemental Fig. 1**.

**c,d**, Reproducibility of the AlphaFold3 prediction of DNA bending for models generated on DNAs of different sequence and/or length. DNA duplexes are superimposed for three AlphaFold3 models in panel **c**, and bend angles from the 25 models shown in **Supplemental Fig. 1** are summarized in panel **d**. Three models for which the DNA was minimally bent are shown in gray. Although the position of the bend along the DNA varies considerably (which causes the pLDDT score of the DNA to be poor in panel **a**), the

predicted arm positions, bend angles, and local deformation of DNA at the bend were all highly reproducible.
**e**, Crystal structure of Topo VI holoenzyme from *Methanosarcina mazei* (pdb: 2q2e)[16].
**f**, Comparison of SPO11–TOP6BL and Top6A–Top6B (pdb: 2q2e) interfaces.
**g,h**, Details of predicted protein-DNA interfaces.
**i**, Correspondence between DNA-contacting residues predicted by the mouse SPO11 AlphaFold3 model and observed in *S. cerevisiae* Spo11 cryo-EM structures[5].

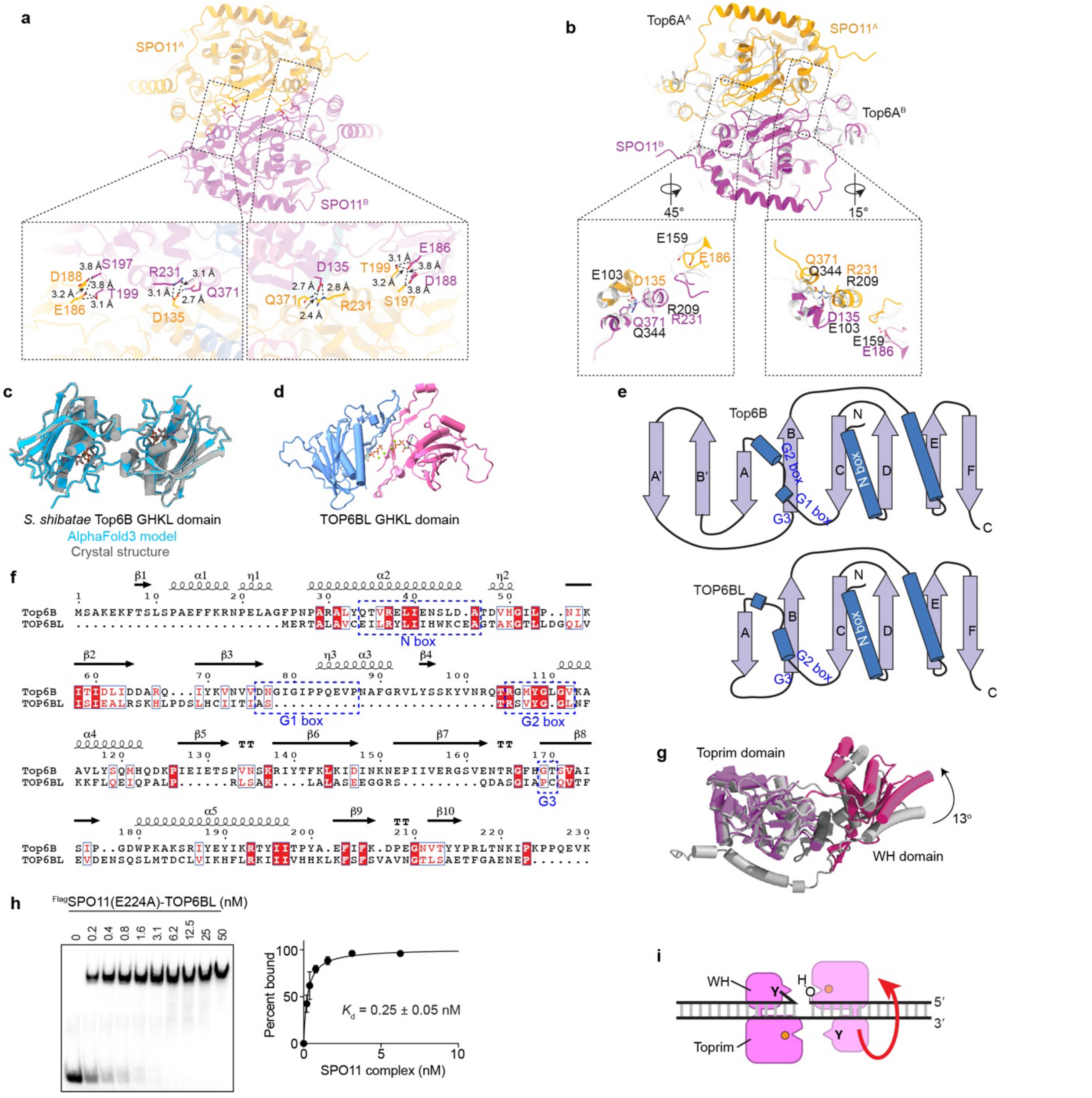


**Extended Data Fig. 4 | Analysis of the SPO11 dimer interface and TOP6BL GHKL domain.**

**a,b**, Details of the SPO11 dimer interface. Panel **a** shows distances between conserved interfacial residues; panel **b** compares the SPO11 dimer interface with that for Top6A in a crystal structure of the *M. mazei* Topo VI holoenzyme (pdb: 2q2e)[16].

**c**, Agreement between AlphaFold3 model and crystal structure (pdb: 1z5a)[37] for an ATP-mediated dimer of the *S. shibatae* Top6B GHKL domain.

**d**, Failure of AlphaFold3 to predict correct ATP-dependent dimerization of the TOP6BL GHKL domain. The dimer interface of the model is in the wrong location (predicted to be on the C-terminus of the GHKL domain), and the ATP molecules are not located at the correct surface between the β sheet and α helices.

**e**, Comparison of topologies of Top6B an TOP6BL GHKL domains. β strands (arrows) and α helices (cylinders) are shown, along with conserved ATP-interacting elements that are highlighted in panel **f.**

**f**, Structure-based sequence alignment between *S. shibatae* Top6B and mouse TOP6BL GHKL domains. A previous alignment based only on the amino acid sequences suggested that TOP6BL has degenerate versions of all three G boxes[11], but alignment based on the AlphaFold3 model shows that the G1 box is missing instead, with degenerate versions of the other two boxes present.

**g**, Hypothetical conformational change between pre-DSB and post-DSB complexes of SPO11 with DNA. A post-cleavage state for mouse SPO11 was modeled by aligning the WH (hot pink) and Toprim (purple) domains separately to the orthologous domains in the yeast Spo11 cryo-EM structure[5]. The AlphaFold3 model (presumptive pre-cleavage state) is shown in gray. This exercise predicts a 13° rotation of the domains relative to one another between the two states.

**h**, EMSA of E224A mutant SPO11 complexes binding to a 5′-labeled 25-bp hairpin substrate with a two-nucleotide 5′ overhang end. A representative gel is shown at left, quantification is at right (mean ± s.d. of n = 3 experiments; apparent $K_d$ given as mean ± s.e.).

**i**, Model for the mechanism of supercoil relaxation by SPO11–TOP6BL complexes. The cartoon (not to scale) shows the predicted arrangement of the WH and Toprim domains of each SPO11 monomer on a nicked DNA molecule, with the 5′ end covalently attached to Y138 of the darker pink monomer on the left, and the 3′ OH bound by the Toprim domain of the lighter colored monomer on the right. The red arrow signifies rotation of the right-hand SPO11 and the DNA arm it is bound to, relative to the left-hand monomer and its bound DNA arm. This rotation, which necessitates disruption of the SPO11 dimer interface, would allow the DNA to swivel around the uncut strand, thereby relaxing supercoils. See **Supplemental Discussion B** for more detail.

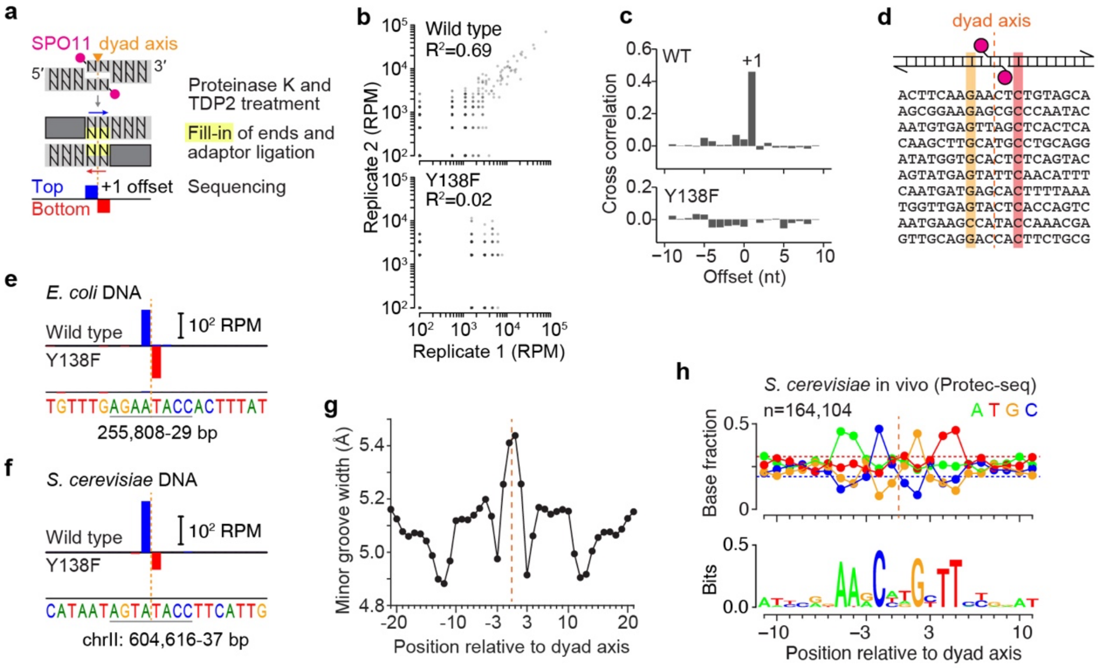


**Extended Data Fig. 5 | Analysis of SPO11 sequence preferences.**
**a**, Schematic description of TDP2-seq.
**b**, Correlation between replicate TDP2-seq datasets for in vitro cleavage of pUC19 with either wild-type (top) or Y138F mutant (bottom) SPO11 complexes. Each point is the read count for a nucleotide position in the plasmid. Positions with no TDP2-seq tags (n = 5,084 and 5,211 for wild type and Y138 mutant maps, respectively) were not used for calculating $R^2$, but were set as $10^{-2}$ for plotting purposes.
**c**, Cross correlation between all top- and bottom-strand reads for in vitro cleavage of pUC19 with wild-type (WT) or Y138F mutant SPO11 complexes.
**d**, Sequence context around the top 10 preferred cleavage sites on pUC19 mapped by TDP2-seq. The bases at –3 and +3 relative to the dyad axis are highlighted.
**e,f**, Examples of preferred in vitro cleavage positions on *E. coli* (**e**) or *S. cerevisiae* (**f**) genomic DNA, presented as in **Fig. 4b** except that read count is given in RPM. Orange dashed lines indicate the inferred dyad axis of cleavage.
**g**, Example of DNA shape properties predicted for the base composition preferred by SPO11. Minor groove width was predicted[70,71] from the mononucleotide frequencies for preferred cleavage sites (n = 5180) on yeast genomic DNA.
**h**, Base composition bias for *S. cerevisiae* Spo11 in vivo. DNA DSB ends generated by Spo11 (n=164,104 sequenced DNA fragments) were identified in Protec-seq maps from *sae2/com1* mutants, in which DSBs remain unresected (data from [44]). Similar base composition biases were also reported for maps of Spo11 oligonucleotides[32,40]. Fractional base composition (above) and sequence logo (below) are shown.

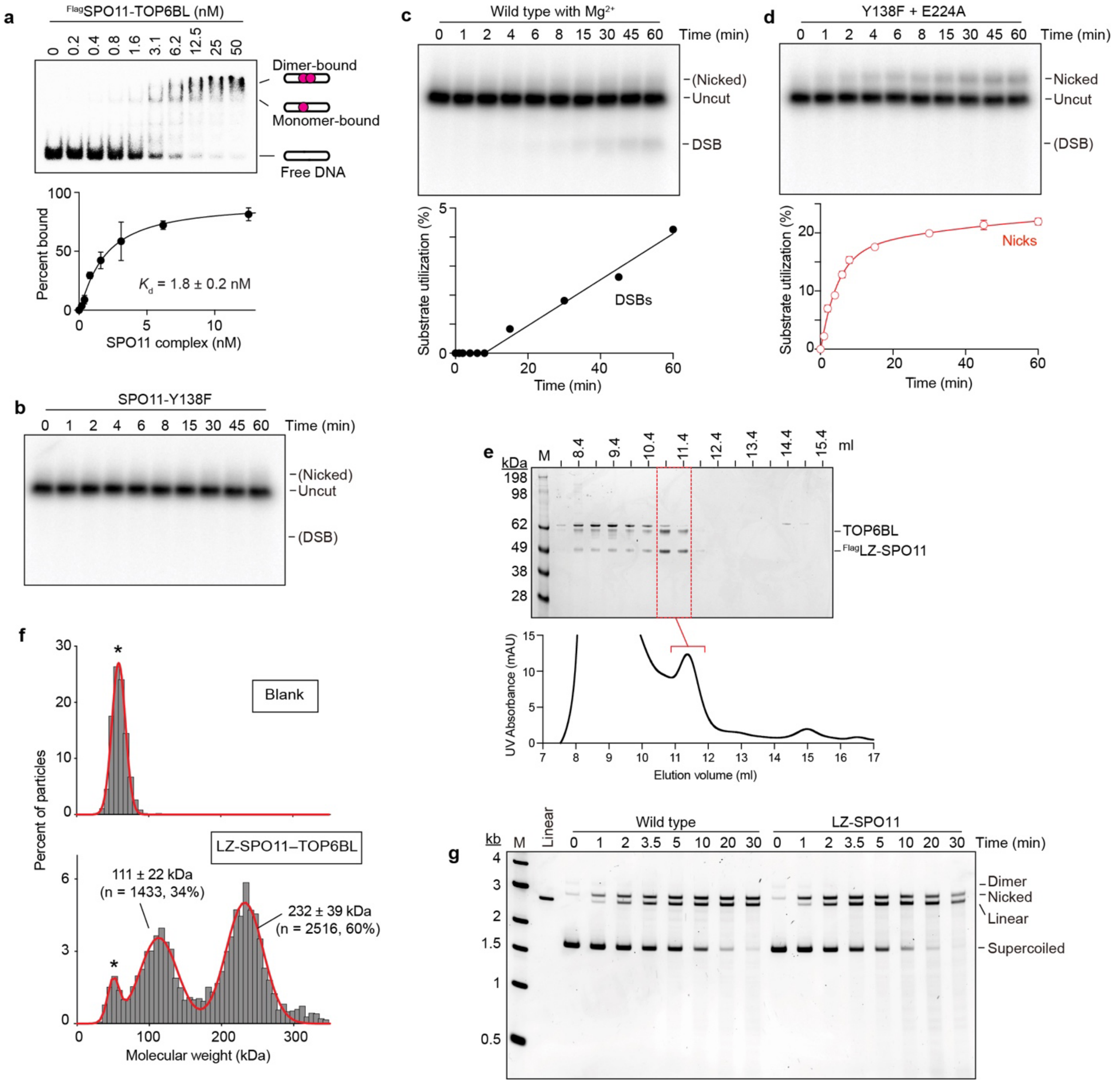


**Extended Data Fig. 6 | Enhancing SPO11 dimerization stimulates DNA cleavage activity.**
**a**, EMSA of SPO11 complexes binding to the $^{32}P$-labeled oligonucleotide substrate shown in **Fig. 4f**. A representative gel is shown above, quantification is below (mean ± s.d. of n = 3 experiments). The apparent $K_d$ (mean ± s.e.) can be viewed as an estimate of the affinity of binding for the first monomer complex.
**b**, Lack of cleavage of the oligonucleotide substrate (0.5 nM) by Y138F mutant SPO11 complexes (10 nM).
**c**, Weak cleavage supported by $Mg^{2+}$. Reactions contained 0.5 nM oligonucleotide substrate, 10 nM wild-type SPO11 complexes, and 5 mM $MgCl_2$. Gel and quantification are provided for a single experiment.
**d**, Nicking-only activity from mixture of Y138F and E224A mutant SPO11 complexes. Reactions contained 0.5 nM oligonucleotide substrate, an equal mixture of mutant protein complexes (10 nM total protein), and 5 mM $MgCl_2$. A representative gel is shown above, quantification is below (mean ± s.d. of n = 3 experiments).
**e**, SEC profile for purification of LZ-SPO11–TOP6BL complexes. Coomassie-stained SDS-PAGE gel (above) and UV profile (below) are shown for chromatography of anti-Flag affinity-purified material.

Fractions pooled for further study are indicated in red. The highlighted peak in the UV chromatogram is at 11.37 ml; the predicted elution volume based on comparison with calibration standards is 11.3 ml.
**f**, Mass photometry analysis of purified LZ-SPO11–TOP6BL complexes. The blank sample (above) lacked protein; protein concentration in the lower graph was 15 nM. Particle counts (gray bars), gaussian density fits (red lines), fitted mean ± s.d., and percentages of total particles are shown. Calculated masses are 115.2 kDa for monomeric and 230.4 kDa for dimeric complexes. Asterisks, background material also present in the blank.
**g**, Representative gel for the experiment quantified in **Fig. 4i**.

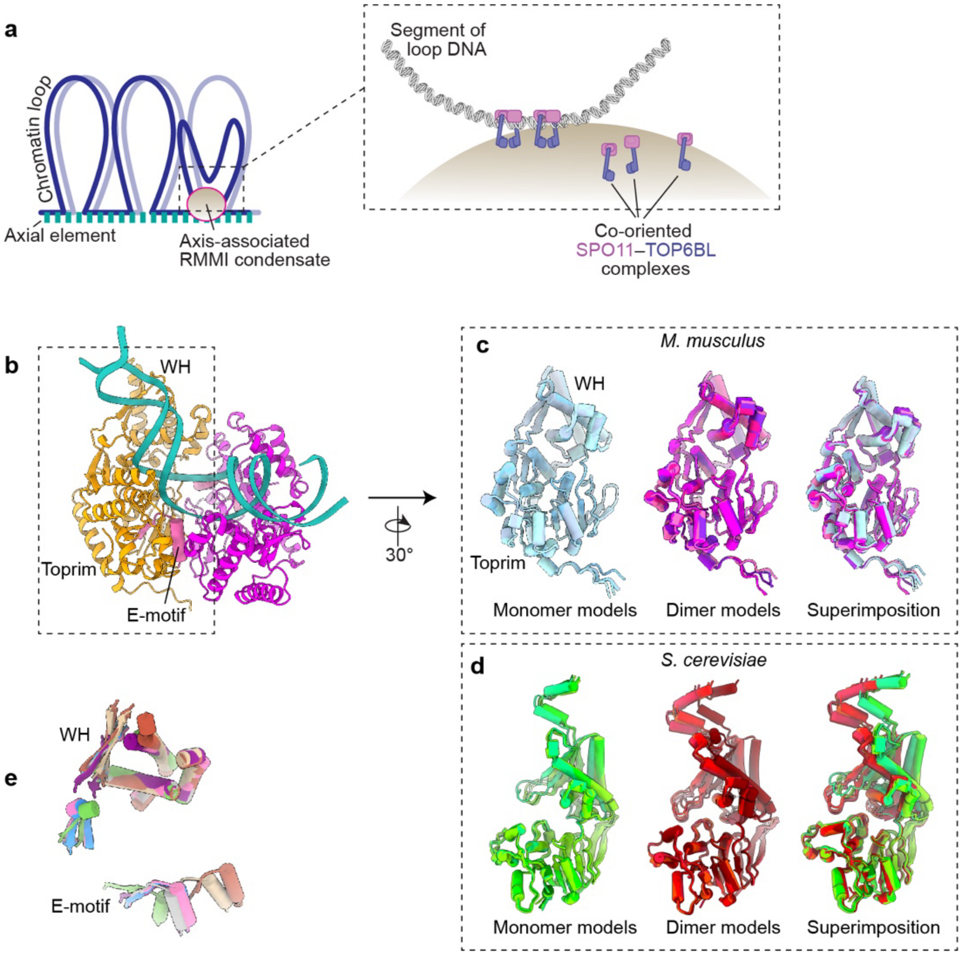


**Extended Data Fig. 7 | A low propensity for dimerization makes SPO11 dependent on accessory factors for activity in vivo.**

**a**, Assembly of the DSB-forming machinery integrated with higher order chromosome structure. The cartoon on the left illustrates the organization of a meiotic chromosome as a linear axial element from which chromatin emanates in loops, with light and dark blue lines representing aligned sister chromatids[9]. In yeast, Rec114, Mei4, and Mer2 proteins assemble cooperatively with DNA and are thought to form axis-associated biomolecular condensates[46]. The mouse orthologs REC114, MEI4, and IHO1 (along with MEI1, which is not found in yeast) likely do so as well (RMMI)[72,73]. The cartoon at right illustrates the hypothesis that these condensates recruit and co-orient clusters of SPO11 core complexes, which can then form dimers to capture and ultimately break a segment of DNA from a nearby chromatin loop[74]. Figure adapted from [5] under a CC-BY license.

**b–d**, SPO11 conformations in monomer and dimer models. Panel **b** presents an overview of the AlphaFold3 prediction of the *M. musculus* SPO11 dimer structure with each SPO11 chain colored as **Fig. 3a**. The E-motif (residues 219–235) from the orange chain is indicated; this segment from the Toprim domain contains the conserved E224 predicted to bind magnesium. Panels **c** and **d** compare the conformations of individual SPO11 chains from AlphaFold3 models of monomers (left, top 5 ranked models) or dimers (middle, top 5 ranked models) for mouse (**c**) or *S. cerevisiae* (**d**). As shown in the superimposed images at right, the monomer and dimer models are similar for the mouse protein but are distinct for yeast, with different relative conformations of the WH domain relative to the Toprim domain. (The dark red dimer model (**d**, middle) predicts an incorrect DNA position, so it is excluded from the monomer-dimer comparison (**d**, right)).

**e**, Comparison of predicted positions of the Toprim domain relative to the WH domain for SPO11 homologs from various species. To evaluate the evolutionary conservation of the distinct monomer conformation in panel **c**, SPO11 monomer structures predicted by AlphaFold3 were compared for *M.*

*musculus* (hot pink), *Homo sapiens* (light blue), *Arabidopsis thaliana* (Spo11-1, purple), *S. cerevisiae* (brown), *Schizosaccharomyces pombe* (tan), and *Sulfolobus shibatae* (Top6A, green). We simplified the display of the WH vs. Toprim domains by aligning on the WH domain (only partial secondary structures of WH used for alignment are shown for simplicity), then showing just the E-motif positions as an indication of the relative position of the Toprim domain. For reference, segments from the mouse SPO11 dimer model are shown in gray. Notably, the monomer structures from *S. cerevisiae* and *S. pombe* exhibit distinct E-motif positions from the other homologs. It is not clear why AlphaFold3 makes different predictions for the yeast protein monomers, but we speculate that it reflects an inherent tendency of the yeast proteins to adopt a conformation that may be less compatible with dimerization.

### Supplemental Discussion

#### A. Evaluating the plausibility of the AlphaFold3 models of dimeric SPO11–TOP6BL complexes bound to DNA

Multiple features of the AlphaFold3 models meet expectations based on empirical structures and functional data for homologous eukaryotic and archaeal proteins. Taken together, these findings suggest that the AlphaFold3 models are plausible representations of dimeric, DNA-bound, pre-DSB complexes.

First, the overall protein architecture closely resembles Topo VI holoenzyme crystal structures[16,17] (**Extended Data Fig. 3e**), with the SPO11–TOP6BL interface in particular matching well with the Top6A–Top6B interface (**Extended Data Fig. 3f**). When individual protein domains are aligned in isolation, the SPO11 WH and Toprim domains align well with crystal structures of archaeal Top6A (rmsd 1.7 Å for WH domains and 2.0 Å for Toprim domains) and cryo-EM structures of yeast Spo11 (rmsd 11.0 Å for WH domains and 7.8 Å for Toprim domains), and the transducer and GHKL domains of TOP6BL align well with archaeal Top6B (rmsd 14 Å for transducer domains and 13 Å for GHKL domains)[5,16,17,19,37]. Moreover, AlphaFold2 models of TOP6BL alone were previously shown to agree well with small-angle x-ray scattering data[31].

Second, the DNA is positioned across a channel formed by the two SPO11 protomers and embraced by the helical transducer domains of the two TOP6BL protomers (**Fig. 3a–c**), as predicted by models of the Topo VI catalytic cycle[16,17,19,38]. Moreover, the modeled DNA binding surface on SPO11 corresponds to the cognate surface of yeast Spo11 defined by hydroxyl radical footprinting and cryo-EM structures[5,6].

Additionally, details of the protein-DNA interface agree well between the AlphaFold3 model and the yeast cryo-EM structures (which are thought to mimic a post-DSB configuration[5]). For example, nearly all of the direct DNA contacts appear to involve SPO11, and are consistent with being primarily interactions with the sugar-phosphate backbone (**Fig. 3d**). Moreover, the AlphaFold3 model places the DNA close to the homologs of many of the SPO11 amino acid residues that engage DNA in the yeast cryo-EM structures[5] (**Fig. 3d and Extended Data Fig. 3g–i**). The similarities include two principally basic patches making backbone contacts (**Extended Data Fig. 3d**); involvement of K175 (yeast K173) in the only clear base contacts (albeit to different positions relative to the dyad axis in yeast and mouse) (**Extended Data Fig. 3h**); and positioning in the minor groove of a series of amino acid side chains from diverse structural elements (previously described as “fingers” in yeast Spo11) (e.g., **Extended Data Fig. 3g**). Notably, however, TOP6BL does appear to contribute directly to DNA binding, particularly around the regions of biased base composition near –10 and +10 positions (**Fig. 3d**), potentially consistent with relatively low affinity binding of purified TOP6BL alone to duplex oligonucleotide substrates ($K_d$ > 500 nM)[31].

Third, two $Mg^{2+}$ ions are positioned within each Toprim domain coordinated by residues E224, D277 and D279 (**Fig. 3e**). This is consistent with expectation for the two-metal-ion catalytic site architecture proposed for type II topoisomerases[75-77]; consistent with structures of Top6A and yeast Spo11[5,19]; and consistent with the importance of these acidic residues for yeast Spo11 function in vivo[78]. Moreover, the metal-binding pocket of each Toprim domain is close to Y138 of the other SPO11 monomer to assemble the expected hybrid active sites, and the two catalytic tyrosines lie close to opposite strands of the DNA as expected (**Fig. 3d,e**). Other aspects of the predicted SPO11 dimer interface also match well with crystal structures for Topo VI (**Extended Data Fig. 4a,b**).

Fourth, the DNA is consistently predicted to be bent (108° ± 21°, mean ± SD of 25 models with DNAs of varying sequence and length) (**Fig. 3b,c and Extended Data Fig. 3c,d**). DNA bending was predicted for Topo VI[38] and was observed for mouse and yeast proteins in AFM experiments (**Fig. 1g**)[6] and in a cryo-EM structure of monomeric yeast core complexes bound to DNA containing a ssDNA gap[5].

**B. Potential mechanism and implications of SPO11 supercoil relaxation activity**

We observe a slow supercoil relaxation activity for SPO11 specifically when nicking has occurred, including conditions where only nicking is possible (e.g., with mixtures of SPO11-Y138F and -E224A mutant proteins). We propose that this topoisomerase activity arises from a swiveling reaction for a dimeric complex stalled at the nicked stage, followed by religation (**Extended Data Fig. 4i**). Specifically, we envision that the weak SPO11 dimer interface allows the two SPO11–TOP6BL complexes to rotate relative to one another, swiveling around the uncut DNA strand opposite to the nick. Reestablishment of the dimer interface after swiveling would then allow the tyrosyl phosphodiester and 3′-OH of the cleaved strand to interact and reseal the nick. Alternatively, swiveling could instead involve release of the noncovalently bound SPO11 monomer, which would necessitate rebinding to effect religation. This scenario could also lead to irreversible nicking. Whether or not SPO11-generated nicks accumulate in vivo has not been definitively established.

In cryo-EM structures, amino acid residues in the yeast Spo11 Toprim domain make direct contacts with the 3′-OH when bound to a DNA end or to a gapped DNA analogous to the product of a nicking reaction[5]. These interactions involve the Toprim domain that would have supported strand cleavage by the catalytic tyrosine on the other Spo11 monomer, i.e., they involve the non-covalently bound Spo11 monomer (**Extended Data Fig. 4i**). Assuming similar contacts for the mouse protein, both SPO11 proteins may remain tightly bound to a nicked DNA: both monomers would have their extensive array of backbone contacts (**Fig. 3d**) supplemented by either covalent phosphotyrosyl attachment for one monomer, or Toprim contacts to the newly generated 3′-OH for the other monomer. If both monomers remain bound, we would further speculate that relaxation activity might be disfavored in vivo because the attachment of SPO11 to nucleoprotein condensates of REC114, MEI4 and other factors would inhibit rotation of SPO11 monomers relative to one another.

Our data also do not establish whether a DSB can be religated. Here and elsewhere[5] we propose that the conformation of both yeast and mouse SPO11 is different in the post-DSB vs. pre-DSB state, at least in part through relative motion of the WH and Toprim domains (**Extended Data Fig. 4g**). Moreover, the post-DSB conformation appears incompatible with dimerization because of steric clashes between the two end-bound complexes[5]. If correct, then a post-DSB conformation would also be incompatible with religation because religation would require reestablishment of a dimer interface similar to the pre-DSB state. In contrast, we suggest that nicking would not allow the covalently bound monomer to relax into the post-DSB configuration without dissociation of the noncovalently bound monomer. This interpretation would allow for nicks to be resealable even if DSBs are not.

**Supplemental Table S1. DNA sequences for binding and cleavage substrates and AlphaFold 3 modeling**

| **a. DNA substrates for EMSAs** | |
|---|---|
| 25 bp hairpin | 5′-TAGCAATGTAATCGTCTATGACGTTAACGTCATAGACGATTACATTGC-3′ |
| 32 bp, top | 5′-TAGCAATGTAATCGTCTATGACGTGTCATAGCGC-3′ |
| 32 bp, bottom | 5′-TAGCGCTATGACACGTCATAGACGATTACATTGC-3′ |
| 31 bp, top | 5′-TAGCAATGTAATCGTCTATGACGTGCATAGCGC-3′ |
| 31 bp, bottom | 5′-TAGCGCTATGCACGTCATAGACGATTACATTGC-3′ |
| 30 bp, top | 5′-TAGCAATGTAATCGTCTATGACGTGATAGCGC-3′ |
| 30 bp, bottom | 5′-TAGCGCTATCACGTCATAGACGATTACATTGC-3′ |
| 29 bp, top | 5′-TAGCAATGTAATCGTCTATGACGTGTAGCGC-3′ |
| 29 bp, bottom | 5′-TAGCGCTACACGTCATAGACGATTACATTGC-3′ |
| 28 bp, top | 5′-TAGCAATGTAATCGTCTATGACGTTAGCGC-3′ |
| 28 bp, bottom | 5′-TAGCGCTAACGTCATAGACGATTACATTGC-3′ |
| 27 bp, top | 5′-TAGCAATGTAATCGTCTATGACGTTAGCG-3′ |
| 27 bp, bottom | 5′-TACGCTAACGTCATAGACGATTACATTGC-3′ |
| 26 bp, top | 5′-TAGCAATGTAATCGTCTATGACGTTAGC-3′ |
| 26 bp, bottom | 5′-TAGCTAACGTCATAGACGATTACATTGC-3′ |
| 25 bp, top | 5′-TAGCAATGTAATCGTCTATGACGTTAG-3′ |
| 25 bp, bottom | 5′-TACTAACGTCATAGACGATTACATTGC-3′ |
| 24 bp, top | 5′-TAGCAATGTAATCGTCTATGACGTTA-3′ |
| 24 bp, bottom | 5′-TATAACGTCATAGACGATTACATTGC-3′ |
| 23 bp, top | 5′-TAGCAATGTAATCGTCTATGACGTT-3′ |
| 23 bp, bottom | 5′-TAAACGTCATAGACGATTACATTGC-3′ |
| 22 bp, top | 5′-TAGCAATGTAATCGTCTATGACGT-3′ |
| 22 bp, bottom | 5′-TAACGTCATAGACGATTACATTGC-3′ |
| 21 bp, top | 5′-TAGCAATGTAATCGTCTATGACG-3′ |
| 21 bp, bottom | 5′-TACGTCATAGACGATTACATTGC-3′ |

| **b. DNA sequences for AlphaFold3 (top strands only)** | |
|---|---|
| 36 bp, seq 1 * | 5′-GTTCGTTGTTACGAAGCATACCCAAACACTTCCCTA-3′ |
| 36 bp, seq 2 | 5′-GCTAGGCAAAAATGGGCATGCCTTTTCTCCATTAAT-3′ |
| 36 bp, seq 3 | 5′-TAGAATCTTGTGTTGGTATACCTCTATATACTAATA-3′ |
| 40 bp | 5′-GTGTTCGTTGTTACGAAGCATACCCAAACACTTCCCTACC-3′ |
| 44 bp | 5′-CGGTGTTCGTTGTTACGAAGCATACCCAAACACTTCCCTACCAC-3′ |

**c. DNA sequence for oligonucleotide cleavage assay****

↓

5′-CGGTTCAAAAGAACCGGAGCTGAATGAAGCC.ATACCAAACGACGAGCGTGACACAAAAGTGTC
ACGCTCGTCGTTTGGT.ATGGCTTCATTCAGCTC-3′

↑

* The top model generated with this DNA was used in the figures (Supplemental File 1).

** A4 loops underlined; central 6 bp around preferred cleavage site from pUC19 in red; arrows, predicted cleavage positions.

**Supplemental Table S2. TDP2-seq mapping statistics**

| Library | No. of reads | No. mapped* | No. uniquely mapped* | No. mapped (mm10)** | No. uniquely mapped (mm10)** |
|---|---|---|---|---|---|
| pUC19 DNA, SPO11-WT, rep 1 | 26,446,950 | 3762 | 3584 | 2,892,802 | 1,212,280 |
| pUC19 DNA, SPO11-WT, rep 2 | 29,036,360 | 4710 | 4544 | 2,580,520 | 1,041,174 |
| pUC19 DNA, SPO11-Y138F, rep 1 | 26,460,313 | 1342 | 1284 | 3,226,714 | 1,375,254 |
| pUC19 DNA, SPO11-Y138F, rep 2 | 24,815,713 | 1246 | 1214 | 2,462,532 | 947,926 |
| *E. coli* DNA, SPO11-WT, rep 1 | 15,211,454 | 409,778 | 393,176 | 7,642,192 | 1,828,402 |
| *E. coli* DNA, SPO11-WT, rep 2 | 25,379,538 | 508,514 | 483,160 | 12,734,598 | 3,129,618 |
| *E. coli* DNA, SPO11-Y138F, rep 1 | 24,380,052 | 472,132 | 449,016 | 3,954,258 | 2,674,612 |
| *E. coli* DNA, SPO11-Y138F, rep 2 | 19,787,344 | 398,098 | 377,752 | 10,068,448 | 2,467,122 |
| *S. cerevisiae* DNA, SPO11-WT, rep 1 | 12,338,811 | 290,444 | 230,234 | 4,744,010 | 1,473,164 |
| *S. cerevisiae* DNA, SPO11-WT, rep 2 | 12,525,951 | 473,424 | 372,610 | 4,470,974 | 1,452,914 |
| *S. cerevisiae*, SPO11-Y138F, rep 1 | 9,191,207 | 231,826 | 171,504 | 4,302,632 | 1,322,228 |
| *S. cerevisiae*, SPO11-Y138F, rep 2 | 7,974,490 | 241,740 | 176,736 | 4,221,372 | 1,295,052 |
| *Mre11-cKO* mouse testis, rep 1 | 93,285,814 | | | 40,041,912 | 10,032,498 |
| *Mre11-cKO* mouse testis, rep 2 | 60,567,142 | | | 24,414,746 | 5,874,994 |

* Reads mapped to pUC19 sequence (pUC19 samples), ASM584v2 genome assembly (*E. coli* samples) or sacCer3 genome assembly (*S. cerevisiae* samples), respectively.

** Reads mapped to mouse mm10 genome assembly. For the in vitro cleavage reactions, these reads arose from mouse cells spiked in as carrier during agarose plug preparation (Methods).

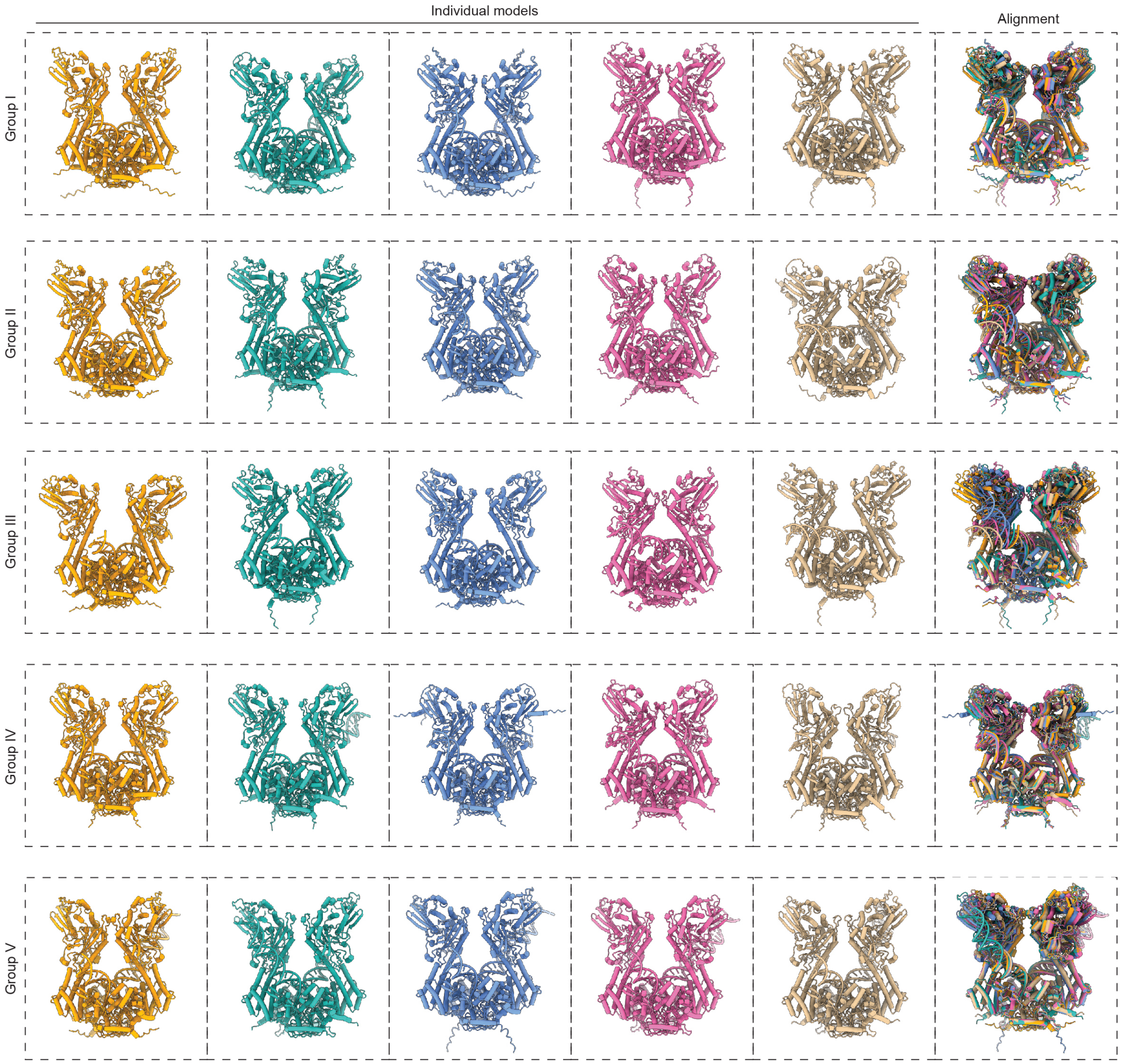


**Suplemental Fig. 1 | Gallery of AlphaFold3 models.** The top five AlphaFold3 models are shown for each of the five DNA sequences shown in Supplemental Table S1b.
